# SpiraMed – A Stereotactic Helix-based Therapy Delivery system for the Human Brain

**DOI:** 10.64898/2026.09.27.754823

**Authors:** Harry Bulstrode, Adam Young, Ifenya Huysman, Dushanth Seevaratnam, King Ki Man, Ho Lim Pak, Grace Wilson, Athena Stamper, Xinyi Wang, Yujun Wang, Xin Tang, An Truong, Saeed Kayhanian, Francisca Ferreira, Mark Plummer, Adam O’Connel, Dominika Bogusiewicz, Daniel Brown, Danny Godfrey, Ronan Daly, George G. Malliaras, David H. Rowitch, Peter J.A. Hutchinson, William P. Gray, Roger A. Barker

## Abstract

Stereotactic needle-based delivery remains the standard for local administration of Advanced Therapy Medicinal Products (ATMPs) to the human brain. ATMP administration typically involves multiple trajectories, presenting cumulative risks and prolonging surgery. Reflux-prone, patchy therapy coverage compromises clinical results. We demonstrate a novel approach, deploying a helical delivery catheter via a single access trajectory per target, referred to as ‘SpiraMed’. Helix retraction is synchronised with therapy delivery, enabling comprehensive target coverage in seconds. Helix pitch and diameter can be precisely tailored to patient-specific target volume and vascular anatomy, as part of the preoperative stereotactic surgical planning process. Testing in agarose phantoms, live sheep and cadaveric human brain confirms enhanced therapy delivery volume, delivery speed and target coverage, with reductions in reflux and predicted risk of bleeding complication. SpiraMed represents a new paradigm, promising to help deliver on the transformative potential of cell and gene therapies across the spectrum of human CNS disease.

## Introduction

Diseases of the Central Nervous System (CNS) affect 3.4 billion people worldwide and account for 11 million deaths per annum^1^. The blood-brain barrier prevents or compromises systemic delivery of many agents^2^, with overall just 8% of candidate CNS therapies succeeding in clinical trials^3,4^. Direct administration to brain parenchyma remains the only approach capable of achieving therapeutic target concentrations of key cell and gene therapy^5–9^, growth factors^10^ and chemotherapy ^11^ combined with acceptable side effect profile^12–15^. This is reflected in year-on-year increases in the number of trials^16^ delivering ATMPs direct to brain: more than 60 gene therapy^17^ and 20 cell replacement ^18^ trials are currently ongoing. All current local ATMP administration is rooted conceptually in stereotactic needle delivery approaches developed pre-clinically in the rodent brain. Established surgical brain delivery technology includes the non CE marked Rehnchrona Legradi device^19^ widely deployed in foundational fetal tissue transplantation trials ^19^, and the CE approved ClearPoint Neuro SmartFlow catheter^20^, commercially licensed for gene therapy in AADC Deficiency and now gaining traction across the spectrum of brain ATMP delivery applications.

Delivery of cells to repair the dopaminergic denervation of the striatum in Parkinson’s disease (PD) is a key exemplar application and subject of 4 recent trial publications^21–24^. However, no device has been developed specifically for delivering such a therapy. In our own TRANSEURO trial (**NCT01898390**)^25^, fetal-derived dopaminergic grafts were administered via 5 straight needle trajectories to each striatum, with multiple deposits per track, in staged operations taking 10-12 hours for each hemisphere. Intracerebral haemorrhage was identified in 2 of 11 patients, reflecting the cumulative risk of multiple needle trajectories. Extended surgical time may have reduced fitness of the cell product, compromising survival, engraftment and outcomes. SpiraMed was conceived in direct response to these concerns, and to address key constraints common to the wider human ATMP delivery landscape.

The human striatum is 3 orders of magnitude greater in volume compared to that of the mouse (35,000mm^3^ vs 23mm^3^)^26^. Hence multiple stereotactic trajectories are typically required, resulting in complex protracted operations^27^. Each trajectory must traverse several centimetres of ‘access’ cortex, incurring direct injury to brain tissue, and a risk of blood vessel injury causing intracerebral haematoma. The latter risk is estimated at up to 1% per needle insertion even with the best pre-operative planning^28^, and in the worst case can result in fatality or long-term disability. Reflux up the needle shaft and out of the target area severely limits infusion pressure and rates, for both conventional and Convection Enhanced Delivery approaches^29^. Small infusate volumes result in small therapy distribution volumes^27^, resulting in patchy target volume coverage and reduced therapy efficacy^27^. Multiple trajectories and slow infusion rates together result in long operations, adding anaesthetic risk and potential to develop pressure sores^30^. Targeting accuracy can suffer as a result of cerebrospinal fluid leak resulting in brain shift^31^. Cell replacement products especially can deteriorate as a result of delays between thawing and administration ^32,33^. Finally long operations threaten the logistics and health economics of future routine clinical application^34^.

Parallel limitations apply to gene therapy delivery in the putamen. Gene therapy diffusion following microinjection is limited to millimetres, resulting in low tissue volume coverage ^35^. Convection Enhanced Delivery (CED)^36^ enhances penetration into brain parenchyma but remains technically demanding: therapy delivery under pressure can exacerbate the tendency to reflux/backflow, necessitating slow therapy delivery and serial re-evaluation with intra-operative MRI. CED of an AAV2-GDNF gene therapy for treatment of Parkinson’s Disease resulted in just 26% putaminal target volume coverage and little improvement in motor scores^37^. CED via posterior/occipital trajectories, aligned with the long axis of the putamen, can improve coverage to over 50% through a single needle track ^38–4027^. However posterior trajectories traverse extensive tracts of eloquent cortex and require prone patient positioning, adding to risk and complexity^28^.

The next generation of brain delivery systems must aim to achieve safer faster comprehensive target volume coverage across a spectrum of ATMP applications. We envisage a single frontal needle access trajectory to each putamen/striatum to minimise risks and duration of surgery. To achieve satisfactory target colume coverage, we propose orthogonal deployment of a delivery catheter spanning the long axis of the putamen target, with scope to tailor catheter dimensions to individual patient target and vascular anatomy. Here we present preclinical proof-of-concept for a novel helix-based delivery solution satisfying these constraints, ‘SpiraMed’.

## Results

### SpiraMed Helix-based Delivery Concept and Prototyping

SpiraMed helix-based delivery entails a single stereotactic trajectory to the brain target using standard neurosurgical stereotactic frame hardware and commercial planning software. A helical delivery catheter is deployed from the SpiraMed needle tip to span the target volume, then retracted, with continuous synchronized therapy delivery into the resulting potential space to minimize reflux (Figure 1A; Supplementary Video 1). Computer aided design and prototype schematics are shown for Generation 1 (mechanically actuated) and Generation 2 (electronically actuated) SpiraMed prototypes (Figure 1B; Supplemental Figures 1-5). Example SpiraMed surgical delivery plans are presented for bilateral ATMP delivery to the putamen via frontal trajectories (Figure 1C), and for chemotherapy delivery to a multifocal high-grade glioma via frontal and parietal trajectories (Figure 1D).

**Figure 1.**
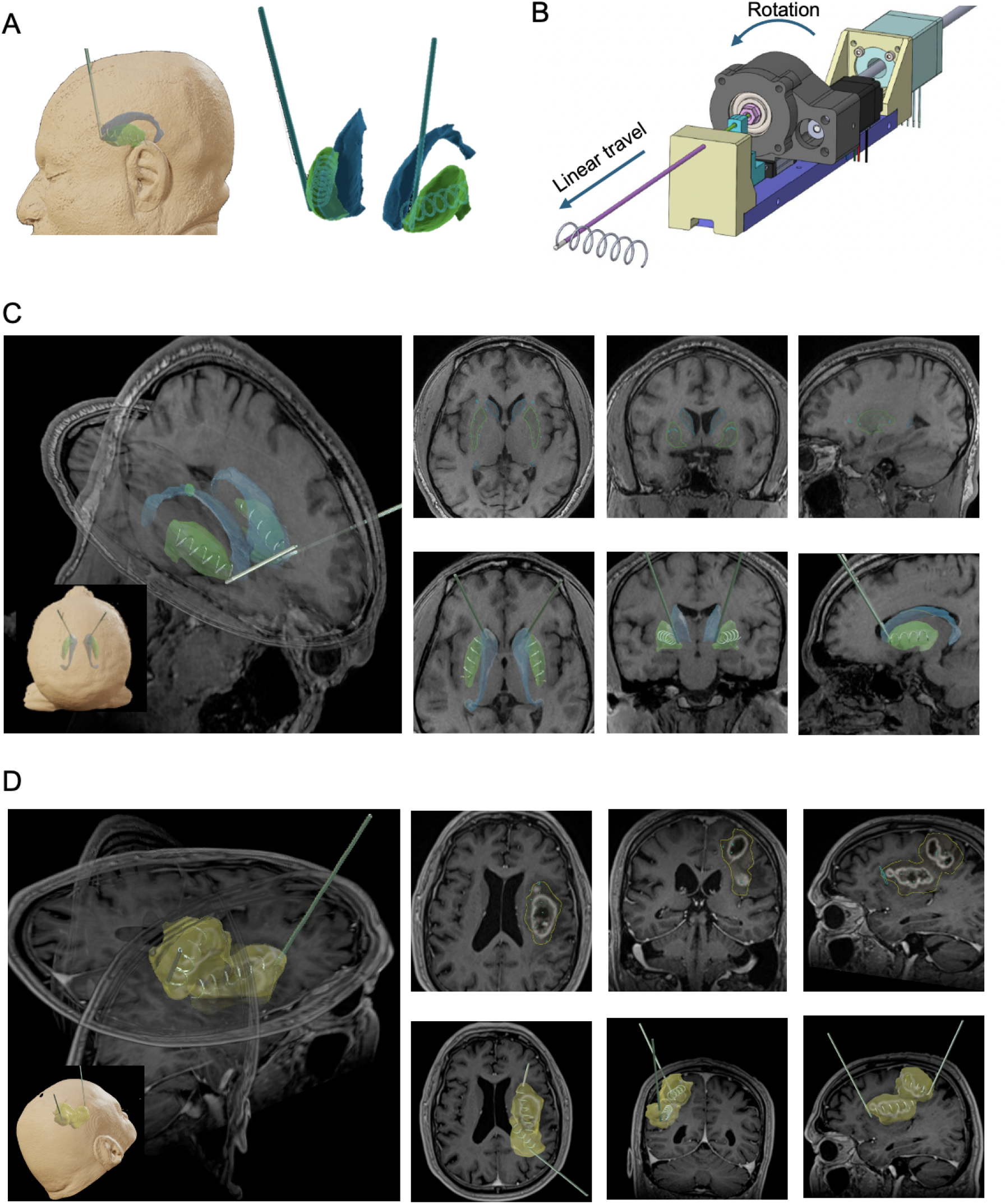
SpiraMed brain therapy delivery concept. **A** Each frontal SpiraMed needle access trajectory permits orthogonal deployment of a SpiraMed delivery catheter spanning the long axis of the striatum **B** SpiraMed Generation 2 Prototype Computer Aided Design illustrating helix deployment **C** Bilateral putaminal delivery strategy envisioned for ATMP delivery in Parkinson’s Disease, AADC deficiency or Huntington’s Disease. Bilateral SpiraMed needle access to the anterior striatum via standard frontal access burrholes is followed by SpiraMed catheter deployment spanning the putamen target volume (green). SpiraMed catheter helix pitch 8mm, diameter 8mm, with 4.5 helix turns is modelled. The trajectory and target nuclei are visualised in single axial, coronal and sagittal planes (upper panels), and as corresponding 3D volumetric overlays (lower panels). **D** Example delivery strategy envisioned for chemotherapy administration to a multifocal high grade glioma, deemed unsuitable for surgical resection. Frontal and parietal SpiraMed approach trajectories are planned, respectively anterior and posterior to primary motor and sensory cortex. SpiraMed catheter helix pitch 8mm, diameter 8mm, with 6 helix turns from the frontal catheter and 4.5 from the parietal is modelled. Trajectories and approximate treatment volumes are visualised in single axial, coronal and sagittal planes (upper panels), and as correspoinding 3D volumetric overlays (lower panels). Surgical plans generated and generated in Brainlab Elements software (Brainlab AG, Munich, Germany).

Generation 1 SpiraMed prototypes employ a hand-wound mechanism to drive the delivery catheter through a helical path within the needle tip, generating fixed helix dimensions (Supplemental Figure 2; detailed discussion Appendix 1). Generation 2 electronic prototypes instead drive the delivery catheter through a planar curved needle tip path, simultaneously applying axial torsion to the catheter to generate the helix (Figure 2A; Supplemental Figures 3,4,5). The latter approach enables each SpiraMed needle tip to deploy a range of helix pitch and diameter combinations, determined by catheter rotation rate □ according to predictions of the Frenet-Serret framework (Figure 2B; detailed discussion Appendix 1). Example helix pitch and diameter values achievable for a needle with tip radius r _tip_=4mm are shown (Figure 2C).

**Figure 2.**
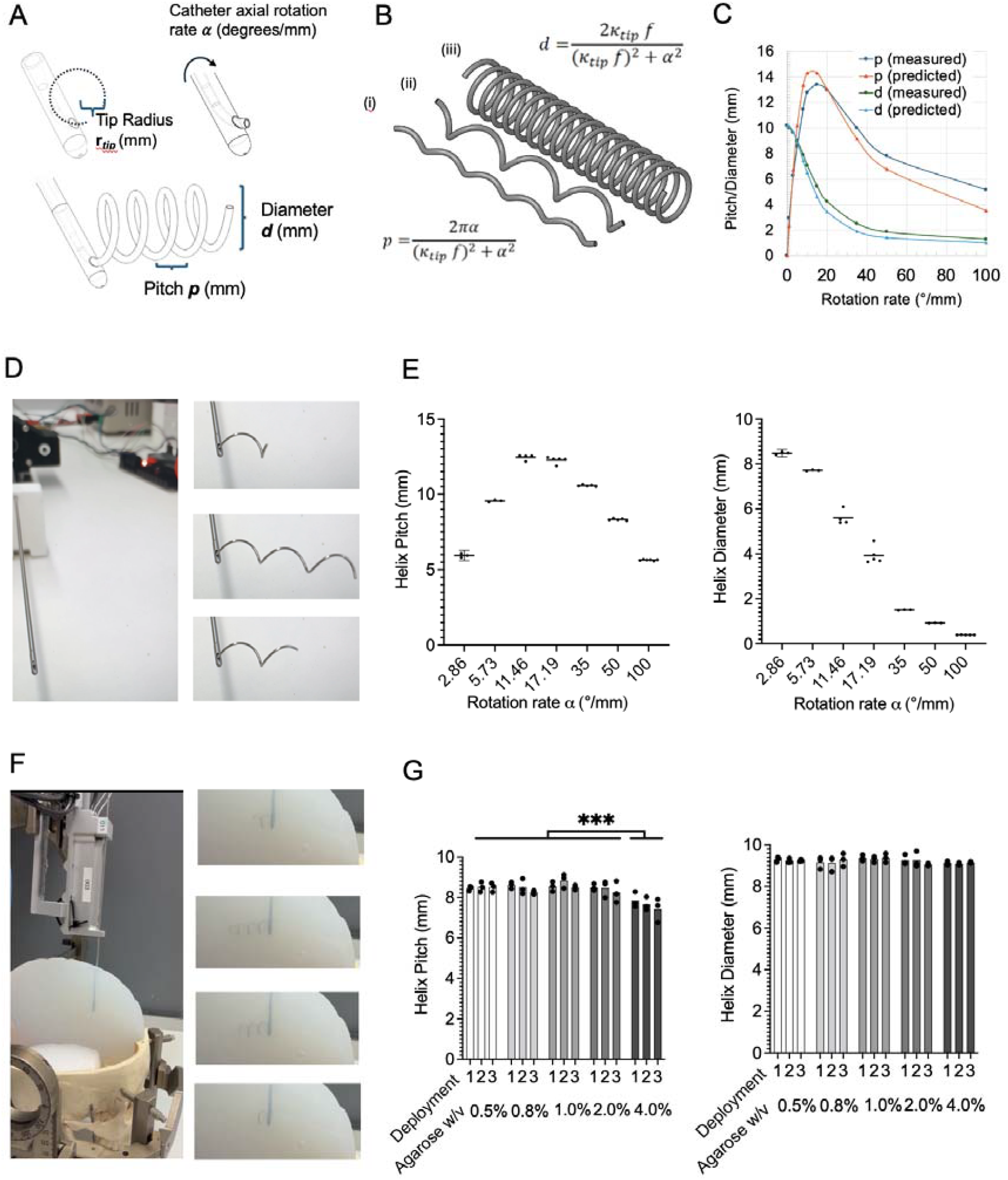
**A** SpiraMed catheter helix pitch **p** and diameter **d** are predicted according to the Frenet-Serret framework as a function of delivery needle internal path radius r _*tip*,_, axial rotation rate ⍰, and constant **f** reflecting the deformation properties of the catheter. **B** The relevant equations and predicted helices are shown for tip radius **r**_*tip*_= 4mm, **f** = 0.86 and catheter rotation rates ⍰ = (i) 35°/mm (ii) 15°/mm (iii) 1°/mm (and see Supplemental Figure 4B) **C** Measured and predicted helix dimensions are shown for delivery needle tip radius **r**_*tip*_= 4mm, **f** = 0.86 as a function of catheter rotation rate (and see Supplemental Figure 4C) **D** SpiraMed benchtop testing example deployment and retraction (**r**_*tip*_=3mm; rotation rate = 5°/mm) **E** At each rotation rate shown, 5 sequential SpiraMed catheter deployments from a single SpiraMed needle tip **r**_*tip*_= 3mm yielded helix pitches (middle panel) and diameters (right panel) shown. **F** SpiraMed benchtop testing in agarose brain phantoms **G** At each agarose concentration shown, three replicate SpiraMed delivery catheters were each deployed and retracted three times through the same SpiraMed needle tip **r**_*tip*_= 3mm, with catheter rotation rate ⍰=5.73°/mm. 2-way ANOVA identifies a significant effect on helix pitch of agarose concentration (F = 12.74, df = 4, p<0.001), but not deployment number (F = 2.05, df = 2, p = 0.1462). Pairwise Tukey’s tests revealed significantly reduced pitch in 4.0% agarose compared to all other conditions (p<0.01 in all cases), with no other comparisons reaching significance. No significant effect on helix diameter of either agarose concentration or deployment number is evidenced.

**Figure 3.**
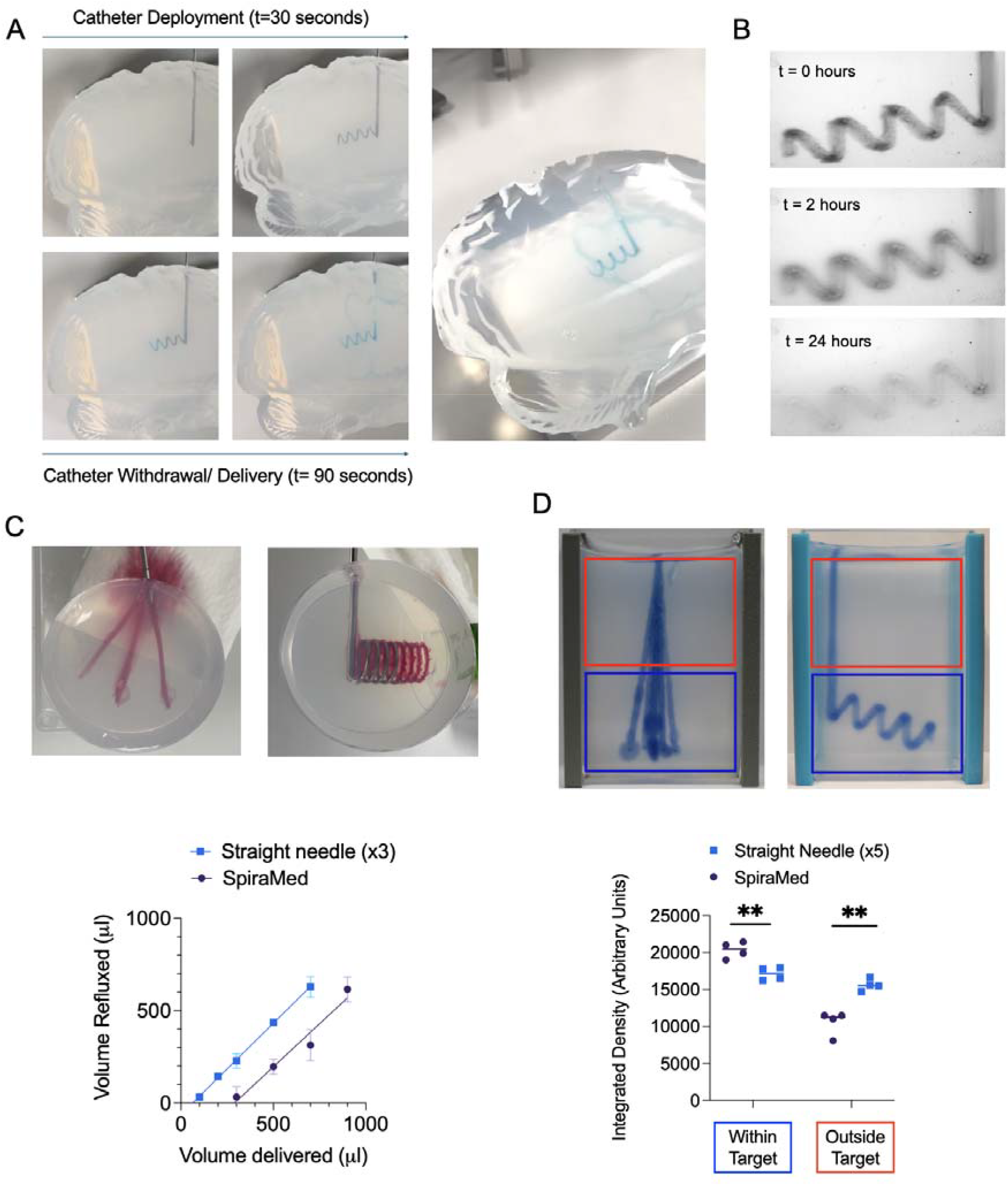
**A** Example SpiraMed catheter deployment and retraction synchronised with reflux-free delivery of dye into 0.6% w/v agarose brain phantom target volumes. **B** ‘Brain-in-a-box’ visualisation of therapy diffusion throughout the target volume over hours following delivery. **C** Dye test volumes (100ul, 200ul, 300ul, 500ul, 700ul, 900ul) were administered in triplicate into fresh agarose blocks, with the volume split between 3 straight needle insertions (left panel) or admininistered in a single SpiraMed delivery (right panel). Volume refluxed was measured by weighing the blotting paper before and after. Estimates of the maximum delivery volume associated with zero reflux (‘reflux threshold’) are provided the x-intercepts of the resulting linear regressions: 64ul (34-90ul) for x3 straight needles, compared to 290ul (181-365ul) for a single Spiramed delivery. F-test confirms significant difference in the xintercepts/ reflux threshold (F-test F=101.0. DFn=1, Dfd=24, P<0.0001). **D** Example images and integrated dye density within (blue ROI)/outside (red ROI) target volume for single SpiraMed (left) and five stereotactic trajectories (right). Total 350 ul methylene blue in aqueous solution was administered, and repeated for 4 replicates in each condition. Significant differences in dye density within target (unpaired t-test t=0.004) and outside target (unpaired t-test t=0.001) are evident.

**Figure 4.**
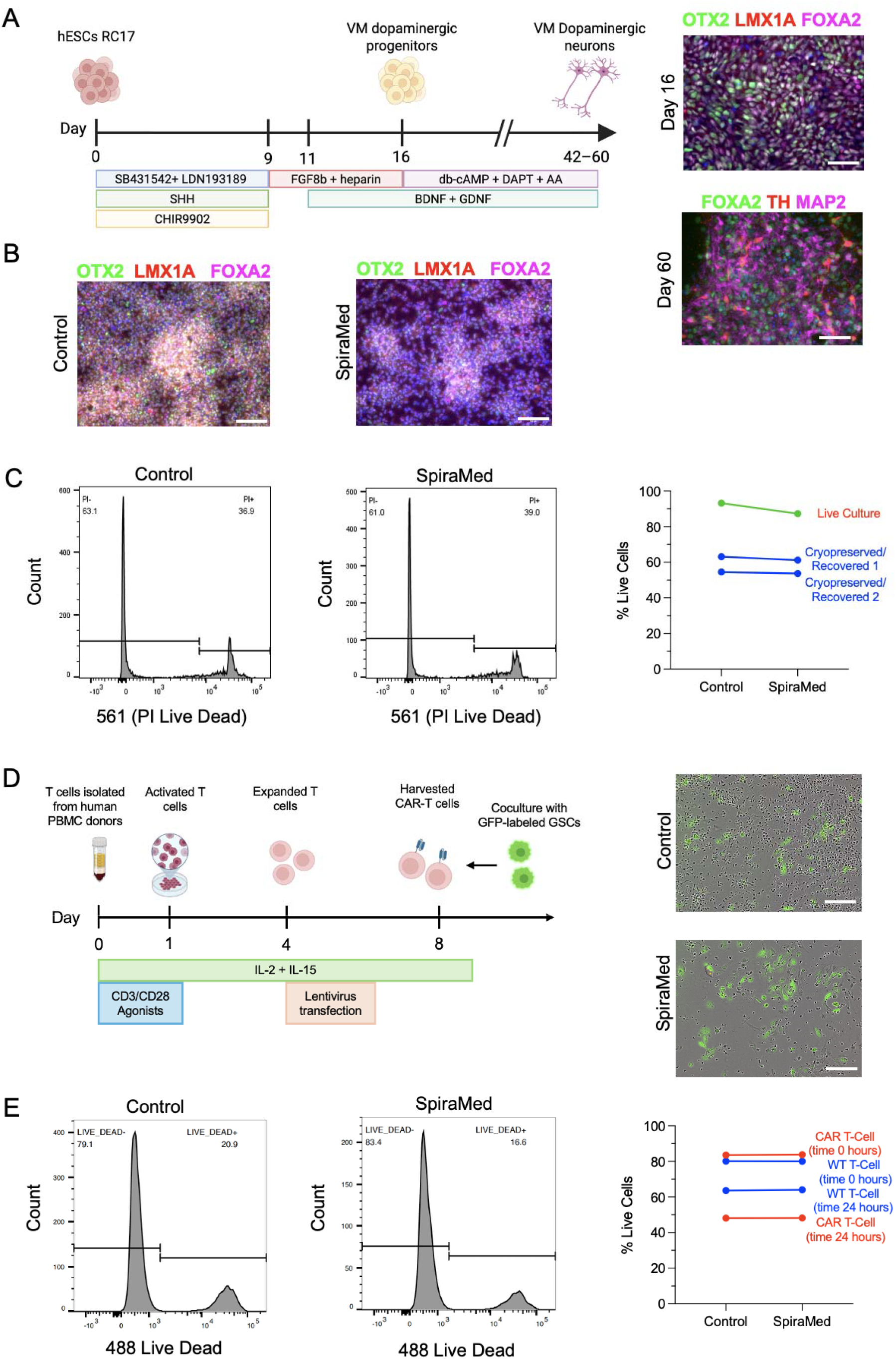
**A:** Differentiation protocol used to generate dopaminergic progenitor cell replacements (left, after Kunath et al.^45^). Progenitor populations suitable for grafting express midbrain markers OTX2, LMX1A, FOXA2 by immunofluorescence at day 16 (upper right panel). Further differentiation *in vitro* yields TH/MAP2 positive dopaminergic neurons by day 60 (lower right panel). Scale bars 20um **B** Immunofluorescence assessment of midbrain progenitor markers in cultured day 16 dopaminergic progenitors (left panel), and after dissociation, aspiration into the SpiraMed delivery catheter, followed catheter deployment and cell delivery back into plates. Scale bar 100um **C** Example flow cytometric assessment of propidium iodide (PI) incorporation (dead cell marker) in cultured mDA progenitors (left) and same following aspiration and delivery through the Spiramed system (middle). Percentage live cells (propidium iodide negative) is shown for 3 biological replicate cell batches (one batch live cell culture to day 16, two batches cryopreserved and recovered 4 hours prior to delivery. No difference in survival between control and SpiraMed-delivered fractions is demonstrated (paired t-test t= 1.9; df= 2; p = 0.20). **D** Diagrammatic representation of protocol used to generate CAR-T cells from primary human donor PBMCs (left) and brightfield images of CAR-T cells cocultured with GFP-labelled glioma stem cells, directly (Control) or following SpiraMed aspiration/delivery. Scale bars 100um **E** Example flow cytometric assessment of 488nm channel dead cell marker incorporation in CAR-T cells (left) and same following aspiration and delivery through the Spiramed system (middle). Percentage live cells (488 negative) post is shown for unmodified T-cell and CAR T-cell batches and paired counterparts, 0 hours and 24 hours following SpiraMed aspiration/delivery. No difference in survival between control and SpiraMed-delivered fractions is demonstrated (t=2.1, df=3, p = 0.13).

**Figure 5.**
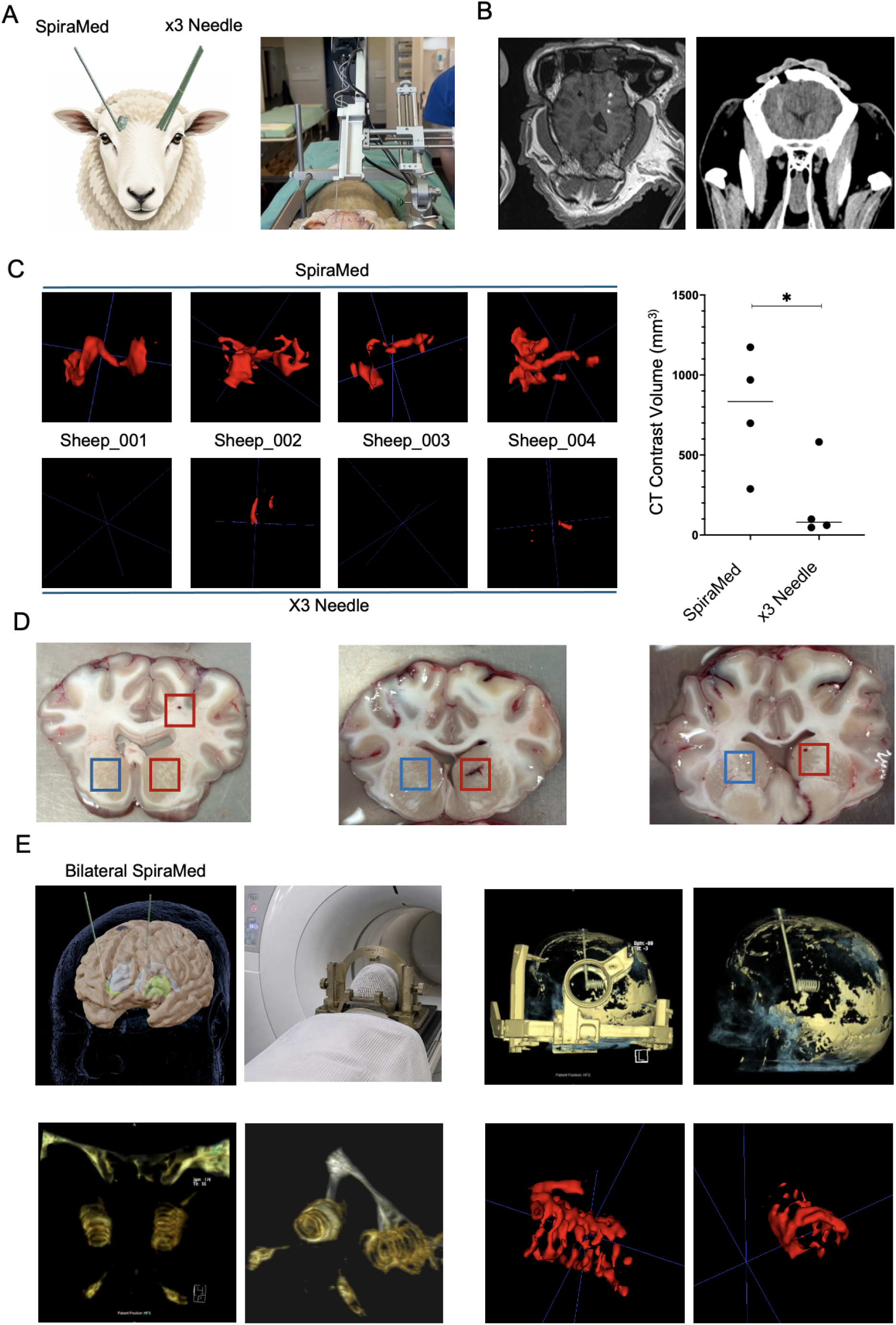
SpiraMed live animal and human cadaver testing. **A** Niopam CT contrast volumes were delivered to the brains of 4 sheep by via frontal SpiraMed needle deployment to one hemisphere, and via 3 SpiraMed needle trajectories without helix deployment to the contralateral hemisphere (left). SpiraMed Generation 2 prototype deployed on the Kopf large animal stereotactic frame ready for catheter deployment and contrast medium delivery is shown (right) **B** Axial MRI demonstrates punctate contrast enhancement coinciding with the helical delivery trajectory (left). Coronal CT imaging demonstrates the burrhole and delivery trajectory achieved in a live sheep (right). **C** Reconstructed 3D contrast volumes for SpiraMed and needle targeted hemispheres in x4 sheep (Unpaired t test with Welch’s correction two-tailed p=0.0497, t=2.539, df =5.25) **D** Gross pathology sections illustrate target volumes following x3 conventional stereotactic trajectories (red) vs single SpiraMed delivery (blue). Microhaemorrhage is also highlighted in the superficial white matter associated with x3 passes. **E** Bilateral putaminal delivery strategy for human cadaveric validation (top left panels). Computed tomography reconstruction demonstrates Leksell G stereotactic frame applied to the cadaveric head, with SpiraMed deployed via a frontal burrhole (top right panels). Contrast delivery to the deep brain in bilateral helical trajectories is evident on CT reconstructions, with skull base bony anatomy also visible (lower left panels), and as volumetric 3D reconstructions (lower right panels; left hemisphere calculated contrast volume 701mm^3^, right hemisphere 994mm^3^).

SpiraMed brain interfacing components are currently realised in 316L surgical stainless steel, ensuring favourable biocompatibility, safety and sustainability, and low material cost. Prototype SpiraMed needles have a 14 gauge (2mm) outside diameter and deploy a 20 gauge catheter (0.91mm outside diameter, 0.6mm inside diameter). Needle and tubing profiles mirror those of existing brain access devices with established safety profiles. SpiraMed helix deployment is pr

### Helix Dimensions and Deployment

Intuitively, helices of approximately equal pitch and diameter appear to represent an acceptable compromise between competing constraints of trajectory length and target coverage, for human putamen and other key brain targets. We chose to target the human putamen (ca 4×2×1cm) using helix pitch and diameter of ∼8mm each (Figure 2D). The dimensions of each replicate helix generated at a given axial rotation rate matched closely, indicating very high *precision* deployment (Figure 2E). Small reproducible systemic *inaccuracies* compared to prediction were evident (Figure 2C; Supplemental Figure 4B), representing parameters not incorporated in our idealised Frenet-Serret framework (e.g. elastic flexion of the delivery needle). These reproducible deviations from theoretical prediction are readily accounted for in any surgical planning software implementation.

We next sought to evaluate potential for helix deflection by the target tissue, deploying and withdrawing each replicate helix repeatedly into agarose gel tissue phantoms (Figure 2F). Agarose blocks of 0.6%-1.0% w/v approximate the consistency of cerebral cortex^41^, while blocks of 2.0% w/v approximate cardiac muscle^42^. No measurable effect of either deployment number or agarose concentration was evident, except in the stiffest 4.0% agarose gels, where helix pitch was fractionally but significantly reduced, whereas helix diameter was unaffected (Figure 2G and Supplemental Figure 2). We conclude that SpiraMed helix deployment is highly predictable at all rotation rates, to a level that comfortably exceeds the 1-1.5mm targeting accuracy acceptable in stereotactic neurosurgery^43^.

### Therapy Delivery in Brain Phantoms

We next evaluated therapy delivery, administering allura red and methylene blue dyes to evaluate therapy coverage and reflux. Dye delivery was synchronised to catheter withdrawal, demonstrating reflux-free delivery of 110ul over a 110mm target trajectory in 90 seconds (Figure 3A). Interval imaging of agarose targets provided qualitative confirmation of rapid consistent target volume coverage by diffusion from the helix trajectory over hours (Figure 3B).

We next sought to estimate SpiraMed volume delivery and reflux compared to a conventional stereotactic needle based approach (Figure 4C). Fixed volumes of dye were injected into an agarose target volume, with each test volume either: (a) divided equally between 3 straight 100mm needle passes during removal of the needle; or (b) delivered through a single SpiraMed trajectory (3mm pitch; 11mm diameter; 8 turns 300mm total catheter deployment). Dye reflux onto blotting paper was quantified by weighing before and after. By performing a linear regression between volume delivered and volume refluxed (Figure 4B right panel), we derived x-intercept values corresponding to a ‘reflux threshold’- the predicted maximum volume that can be delivered via each approach prior to onset of reflux. The cumulative reflux threshold for delivery across 3 straight needle passes was 64.0µl (95% CI 34.3-90.2µl) whereas the threshold for a single SpiraMed delivery was 290.5µl (95% CI 181.1-365.1µl).

Finally we sought to compare SpiraMed target volume coverage compared to conventional stereotactic delivery, by means of 4 replicate deliveries of aqueous methylene blue solution into agarose, with optical quantification of retention within the arbitrary cuboid target volume vs that which refluxed out of this volume (Figure 3D). The mean integrated density within the target volume for a single SpiraMed delivery was 20351 compared to 17136 for x5 conventional stereotactic deliveries (t=4.53, q = 0.004). The mean integrated density outside the target volume (representing reflux into ‘access cortex) was 10517 for SpiraMed compared to 15603 for x5 conventional stereotactic deliveries (t=5.56, q=0.0029).

We conclude that SpiraMed offers significantly enhanced therapy delivery to target compared to conventional stereotactic delivery, with reduced reflux.

### ATMP Biocompatibility

The 316L surgical steel alloys employed in SpiraMed construction are considered to demonstrate good short-term biocompatibility with brain tissue and ATMP therapy classes^19,44^. However we considered that helix deployment could conceivably result in heat or fluid dynamic insult to ATMP products. We opted to test for impact on cell therapies which are arguably the most exacting use case.

We firstly generated fresh and cryopreserved ventral midbrain dopaminergic (vmDA) progenitor cell batches, equivalent to cell products currently in clinical trials ^21–24^ (Figure 4A and Online Methods)^45^. Quality control by immunofluorescence and quantitative polymerase chain reaction confirmed robust midbrain dopaminergic progenitor identity, and mature dopaminergic neuron differentiation potential (Figure 4A; Supplementary Figure 6). A suspension of vmDA progenitors (8×10 ^6^ cells/ml) was aspirated into the SpiraMed needle/catheter assembly. 5 helix turns were deployed, then retracted at 1mm/second with simultaneous delivery of cell suspension at 1ul/second into a fresh microtube. Live cultured and recovered crypopreserved progenitors demonstrated equivalent viability by flow cytometry with and without SpiraMed delivery, and re-plating confirmed preservation of marker expression (Figure 4B, 4C).

Secondly, we generated Chimeric Antigen Receptor T-cell batches using peripheral blood mononuclear cell (PBMC) isolates from healthy donors: intra-tumoural delivery of Chimeric Antigen Receptor T-cells is a key emerging therapy in children especially^46,47^. We assayed CAR-T activity in coculture with GFP-labelled glioma stem cells (Figure 4D). Flow cytometry indicated no differences in viability between SpiraMed-delivered and control T-cells (unmodified and CAR-T bearing) at either 0 hours or 24 hours post delivery (Figure 4E).

### Large animal *in vivo* and human cadaveric testing

In order to confirm safety of deployment and satisfactory therapy delivery *in vivo*, we undertook stereotactic delivery of contrast medium to deep brain targets in sheep in Adelaide, Australia. In each animal, equal volumes of contrast medium were delivered via SpiraMed to one hemisphere, and via 3 conventional stereotactic trajectories to the contralateral hemisphere (Figure 5A, B). Contrast volumes retained in the deep brain target were quantified by volumetric MRI reconstruction (Figure 5C), with significantly greater retention demonstrated using SpiraMed compared to conventional trajectories (mean 782.1ul vs 196.8ul; Welch-corrected t=2.539, df=5.25, p=0.0497). No haemorrhagic complications were evident in any SpiraMed-targeted hemisphere, whereas microhaemorrhages were evident in the target region and in superficial access cortex in one conventionally-targeted hemisphere (Figure 5D).

Finally, we sought to evaluate integration of SpiraMed into a standard stereotactic neurosurgical workflow. To do this we secured cadaveric human heads in a Leksell G stereotactic frame. We scanned the heads in a GE CT scanner to register the deep brain targets in stereotactic coordinate space, and employed pre-planned trajectories to the anterior putamen on each side of the brain (Figure 5E). Following SpiraMed deployment, 1000ul of radio-opaque contrast medium was administered to each side of the brain by repeated bolus push interspersed with catheter retraction over a total of one minute. Post-procedure CT scan demonstrated satisfactory volume coverage in the expected helical configuration bilaterally (Left 700.8mm^3^ Right 994.4mm^3^; Figure 7D, E, F).

## Discussion

SpiraMed represents a novel approach to brain delivery, conceived to address inherent limitations in current technology. Our approach has been informed by Radially Branched Deployment (RBD)^48^, also developed to enhance target volume coverage. RBD however requires repeated deployment, retraction and reorientation of the delivery catheter, resulting in a complex multi-step surgical workflow. RBD remains prone to reflux and has not yet progressed beyond preclinical evaluation. In contrast SpiraMed offers rapid ‘single-pass’ therapy delivery, improving therapy volume delivered, target volume coverage, reflux and predicted surgical safety (summarized Table 1).

**Table 1.** Comparison of key practical surgical and delivery parameters, experimentally-determined or predicted for SpiraMed vs conventional stereotactic delivery. Predicted target volume (V =pr^2^h) is based on the assumption of effective therapy delivery circumferentially 3mm around each delivery trajectory. Predicted ICH risk is based on an estimated risk of haemorrhage of 0.1% per 1cm brain trajectory, and total trajectory distances as shown.

|  | <u>SpiraMed</u> | <u>Conventional Stereotactic Delivery</u> |
| --- | --- | --- |
| **Surgery Format** | Single pass per hemisphere | 3 passes per hemisphere |
| **Surgery Duration**<br>(Bilateral) | **2-3 hours** | **4-6 hours** |
| **Reflux Threshold**<br>(see Figure 3c) | **290ul**<br>(Single pass) | **64ul**<br>(3 passes) |
| **Target coverage**<br>(Predicted see legend) | **3.7cm<sup>3</sup>**<br>(13cm trajectory within target) | **1.3 cm<sup>3</sup>**<br>(3 trajectories, each 1.5cm within target) |
| **Trajectory Distance**<br>(Per hemisphere) | **21 cm**<br>8cm (needle) + 13cm (helix) | **24cm**<br>8cm (needle) x 3 passes |
| **Predicted ICH risk**<br>(Bilateral surgery) | **4.2%** | **4.8%** |

SpiraMed needle access and catheter deployment entails a ∼50% reduction in total trajectory length through brain tissue compared to five conventional stereotactic trajectories, potentially conferring a corresponding reduction in tissue injury, and especially risk of a bleeding complication due to blood vessel damage. We acknowledge that this simple metric fails to account for the high vascularity of the putamen itself, and for the clinical significance of haemorrhage in this region. Additional targeted vascular safety studies will be required to support future patient application.

Reductions in surgery duration and simplified workflows achievable using SpiraMed delivery could be critical to the scale up of brain ATMP delivery to routine clinical application, and could also enable new clinical trial paradigms. For example, patients with Parkinson’s Disease eligible for subthalamic nucleus deep brain stimulation (STN DBS) could be offered blinded randomisation to DBS alone, or DBS and SpiraMed dopaminergic cell replacement under the same anaesthetic. This could address outstanding logistical and ethical barriers around blinding and sham surgery in this field^49^. ‘Adaptive’ DBS hardware sampling of STN local field potentials could also help gauge cell replacement functional engraftment, which is not well assessed in current trials.

SpiraMed was conceived to facilitate delivery of cell replacements. However we envisage that in combination with Convection Enhanced Delivery it could also facilitate rapid delivery of large volume gene therapy infusates, enhancing therapy volume of distribution while obviating the need for posterior trajectories. Refinements in catheter design, together with integration of delivery pump and realtime pressure/flow sensing technology, will be key to achieving this. Additional areas of focus in developing the next generation of ‘patient-ready’ SpiraMed hardware will include robust stereotactic surgical planning software implementation, and a failsafe catheter retraction mechanism – in the event of power failure for example. Implementation of SpiraMed technology in titanium alloys may ultimately be desirable for application in the context of intraoperative Magnetic Resonance Imaging, and preliminary trials of titanium needle tips and catheters have presented no unexpected technical concerns. Finally, it has not escaped our notice that the specifics of helix-based delivery immediately suggest integration into robotic surgical delivery platforms, to enable still faster more straightforward ATMP delivery surgical workflows.

SpiraMed technology offers the promise of safer, faster, and more effective direct brain delivery, and could help realise the potential of emerging ATMP classes across the spectrum of CNS disease.

## Supporting information

Appendix 1 - Frennet Serret Framework

## Author Contributions

**HB** devised the SpiraMed helical delivery concept, secured funding, directed or performed all experimental validation and surgical planning work, and drafted the manuscript. **AY** helped develop the SpiraMed concept, secured funding, and led *in vivo* experimental work. **IH** engineered SpiraMed helix deployment including hardware and software design, building and testing of Generation 1 and Generation 2 SpiraMed prototypes. **KKM, KP, DS** performed experimental work. **GW, AS, XW, YW, XT, AT and SK** contributed to cell culture and flow cytometry experiments. **FF** performed imaging volume reconstructions and quantification. **DB** and **DB** oversaw human cadaveric work. **MP** and **AO** oversaw *in vivo* work. **DG** led the eg technology engineering team. **RD and GM** oversaw academic engineering testing. **DHR, WPG, PJAH, and RAB** advised on development and testing and contributed academic oversight and advice on regulatory and translational aspects.

## Acknowledgements

The authors are deeply indebted to the Cambridge Enterprise team - Tom Mentlak, Katja Kostelnik, Rachel Atfield and Hannah Pape with respect to funding and intellectual property protection. UK priority patent application GB2412772.2 was filed in August 2024 by Cambridge Enterprise (CE, on behalf of University of Cambridge), including device and method for the formation of a helical coil across varied applications. The filing entered PCT (PCT/GB2025/051907) in August 2025, and is now published (WO2026/047355). The filing will enter the national phase in February 2027 in territories to include all potential main markets and manufacturing locations. CE is responsible for commercialisation of SpiraMed, has performed in depth prior art and patentability searches and market research analyses, and will continue to explore new IP opportunities arising during the project. All future IP arising from the proposed work will be owned by the University of Cambridge and managed through Cambridge Enterprise.

## References

1. Organization, W. H. Global Status Report on Neurology. https://www.who.int/publications/i/item/9789240116139 (2025).

2. Wu, D. et al. The blood–brain barrier: Structure, regulation and drug delivery. Signal Transduct. Target. Ther. 8, 217 (2023).

3. Miller, G. Is Pharma Running Out of Brainy Ideas? Science 329, 502–504 (2010).

4. Bhunia, S. et al. Drug Delivery to the Brain: Recent Advances and Unmet Challenges. Pharmaceutics 15, 2658 (2023).

5. Aboody, K., Capela, A., Niazi, N., Stern, J. H. & Temple, S. Translating Stem Cell Studies to the Clinic for CNS Repair: Current State of the Art and the Need for a Rosetta Stone. Neuron 70, 597–613 (2011).

6. Lindvall, O. & Kokaia, Z. Stem cells in human neurodegenerative disorders — time for clinical translation? J. Clin. Investig. 120, 29–40 (2010).

7. Axelsen, T. M. & Woldbye, D. P. D. Gene Therapy for Parkinson’s Disease, An Update. J. Park.’s Dis. Preprint, 1–21 (2018).

8. Stamper, A., Bulstrode, H. & Barker, R. A. Stem cell treatments and Parkinson’s disease: Science and misconceptions. J. Park.s Dis. 1877718x261434672 (2026) doi:10.1177/1877718x261434672.

9. Klein, R. L. et al. NGF gene transfer to intrinsic basal forebrain neurons increases cholinergic cell size and protects from age-related, spatial memory deficits in middle-aged rats. Brain Res. 875, 144–151 (2000).

10. Buttery, P. C. & Barker, R. A. Gene and Cell-Based Therapies for Parkinson’s Disease: Where Are We? Neurotherapeutics 17, 1539–1562 (2020).

11. Sawyer, A. J., Piepmeier, J. M. & Saltzman, W. M. New methods for direct delivery of chemotherapy for treating brain tumors. Yale J. Biol. Med. 79, 141–52 (2006).

12. Bruce, J. N. et al. Regression of Recurrent Malignant Gliomas With Convection-Enhanced Delivery of Topotecan. Neurosurgery 69, 1272–1280 (2011).

13. D’Amico, R. S., Aghi, M. K., Vogelbaum, M. A. & Bruce, J. N. Convection-enhanced drug delivery for glioblastoma: a review. J. Neuro-Oncol. 151, 415–427 (2021).

14. Osorio, M. J. et al. Concise Review: Stem Cell□Based Treatment of Pelizaeus□Merzbacher Disease. STEM CELLS 35, 311–315 (2017).

15. Li, H. et al. Gene suppressing therapy for Pelizaeus-Merzbacher disease using artificial microRNA. JCI Insight 4, e125052 (2019).

16. Elder, J. B. et al. Direct delivery of gene- and cell-based therapies to the nervous system. Image-Guided Biologic Therapies: Neurosurgeons Innovating Treatment Excellence Summit summary. J. Neurosurg. 143, 1431–1441 (2025).

17. Patel, R. V., Nanda, P. & Richardson, R. M. Neurosurgical gene therapy for central nervous system diseases. Neurotherapeutics 21, e00434 (2024).

18. Kirkeby, A., Main, H. & Carpenter, M. Pluripotent stem-cell-derived therapies in clinical trial: A 2025 update. Cell Stem Cell 32, 10–37 (2025).

19. Lindvall, O. et al. Human Fetal Dopamine Neurons Grafted Into the Striatum in Two Patients With Severe Parkinson’s Disease: A Detailed Account of Methodology and a 6-Month Follow-up. Arch. Neurol. 46, 615–631 (1989).

20. Han, S. J., Bankiewicz, K., Butowski, N. A., Larson, P. S. & Aghi, M. K. Interventional MRI-guided catheter placement and real time drug delivery to the central nervous system. Expert Rev. Neurother. 16, 635–639 (2016).

21. Tabar, V. et al. Phase I trial of hES cell-derived dopaminergic neurons for Parkinson’s disease. Nature 641, 978–983 (2025).

22. Sawamoto, N. et al. Phase I/II trial of iPS-cell-derived dopaminergic cells for Parkinson’s disease. Nature 641, 971–977 (2025).

23. Chang, J. W. et al. Phase 1/2a clinical trial of hESC-derived dopamine progenitors in Parkinson’s disease. Cell 10.1016/j.cell.2025.09.010 (2025) doi:10.1016/j.cell.2025.09.010.

24. Paul, G. et al. Human embryonic stem cell-derived dopaminergic cells for Parkinson’s disease: a phase 1/2 open-label trial. Nat. Med. 1–9 (2026) doi:10.1038/s41591-026-04525-0.

25. Barker, R. A. et al. The TransEuro open-label trial of human fetal ventral mesencephalic transplantation in patients with moderate Parkinson’s disease. Nat. Biotechnol. 1–9 (2025) doi:10.1038/s41587-025-02567-2.

26. Kozlov, A. et al. Mouse and human striatal projection neurons compared-somatodendritic arbor, spines and in silico analyses. PLOS Comput. Biol. 21, e1013569 (2025).

27. Richardson, R. M. et al. Data-driven evolution of neurosurgical gene therapy delivery in Parkinson’s disease. J. Neurol., Neurosurg. Psychiatry 91, 1210–1218 (2020).

28. Maheshwari, S. et al. Dopaminergic Cell Replacement for Parkinson’s Disease: Addressing the Intracranial Delivery Hurdle. J. Park.’s Dis. 14, 415–435 (2024).

29. Mehta, A. M., Sonabend, A. M. & Bruce, J. N. Convection-Enhanced Delivery. Neurotherapeutics 14, 358–371 (2017).

30. Cheng, H. et al. Prolonged operative duration is associated with complications: a systematic review and meta-analysis. J. Surg. Res. 229, 134–144 (2018).

31. Chapelle, F. et al. Early Deformation of Deep Brain Stimulation Electrodes Following Surgical Implantation: Intracranial, Brain, and Electrode Mechanics. Front. Bioeng. Biotechnol. 9, 657875 (2021).

32. Hakami, A. et al. Graft ischemia post cell transplantation to the brain: Glucose deprivation as the primary driver of rapid cell death. Neurotherapeutics 22, e00518 (2025).

33. Kim, T. W. et al. TNF-NF-κB-p53 axis restricts i n vivo survival of hPSC-derived dopamine neurons. Cell 187, 3671–3689.e23 (2024).

34. Zhang, R. et al. Eladocagene Exuparvovec for the Treatment of Aromatic l-Amino Acid Decarboxylase Deficiency (AADCd): An Economic Evaluation from a US Perspective. PharmacoEconomics 44, 25–41 (2026).

35. Chu, Y., Bartus, R. T., Manfredsson, F. P., Olanow, C. W. & Kordower, J. H. Long-term post-mortem studies following neurturin gene therapy in patients with advanced Parkinson’s disease. Brain 143, 960–975 (2020).

36. Bobo, R. H. et al. Convection-enhanced delivery of macromolecules in the brain. Proc. Natl. Acad. Sci. 91, 2076–2080 (1994).

37. Heiss, J. D. et al. Trial of magnetic resonance–guided putaminal gene therapy for advanced Parkinson’s disease. Mov. Disord. 34, 1073–1078 (2019).

38. Bankiewicz, K. S. et al. AAV viral vector delivery to the brain by shape-conforming MR-guided infusions. J. Control. Release 240, 434–442 (2016).

39. Laar, A. D. V. et al. Intraputaminal Delivery of Adeno□Associated Virus Serotype 2–Glial Cell Line–Derived Neurotrophic Factor in Mild or Moderate Parkinson’s Disease. Mov. Disord. 40, 1297–1306 (2025).

40. Sudhakar, V. et al. Infuse-as-you-go convective delivery to enhance coverage of elongated brain targets: technical note. J. Neurosurg. 133, 530–537 (2020).

41. Chen, Z.-J. et al. A realistic brain tissue phantom for intraparenchymal infusion studies. J. Neurosurg. 101, 314–322 (2004).

42. Tejo-Otero, A. et al. Soft-Tissue-Mimicking Using Hydrogels for the Development of Phantoms. Gels 8, 40 (2022).

43. Rajabian, A. et al. Accuracy, precision, and safety of stereotactic, frame-based, intraoperative MRI-guided and MRI-verified deep brain stimulation in 650 consecutive procedures. J. Neurosurg. 138, 1702–1711 (2023).

44. Sargioti, N., Levingstone, T. J., O’Cearbhaill, E. D., McCarthy, H. O. & Dunne, N. J. Metallic Microneedles for Transdermal Drug Delivery: Applications, Fabrication Techniques and the Effect of Geometrical Characteristics. Bioengineering 10, 24 (2022).

45. Kunath, T. et al. Midbrain dopaminergic differentiation of human pluripotent stem cells v1. 10.17504/protocols.io.bddpi25n (2020) doi:10.17504/protocols.io.bddpi25n.

46. Baldo, G. D. et al. The peculiar challenge of bringing CAR-T cells into the brain: Perspectives in the clinical application to the treatment of pediatric central nervous system tumors. Front. Immunol. 14, 1142597 (2023).

47. Chan, J. T. N., Henley-Waters, J. & Kayhanian, S. Chimeric antigen receptor (CAR)-T-cell therapy for glioblastoma: what can we learn from the early clinical trials? A systematic review. Neuro-Oncol. Adv. 7, vdaf115 (2025).

48. Silvestrini, M. T. et al. Radially Branched Deployment for More Efficient Cell Transplantation at the Scale of the Human Brain. Stereotact. Funct. Neurosurg. 91, 92–103 (2013).

49. Tabar, V. & Barker, R. A. Sham surgery for the trialing of cell-based therapies to the CNS may not be necessary. Cell Stem Cell 31, 158–160 (2024).

