## Supplementary material for "SpiraMed – A Stereotactic Helix-based Therapy Delivery system for the Human Brain": Online Methods and Supplemental Figures

**SpiraMed prototypes**

Generation 1 and 2 SpiraMed technology were designed and built by eg technology according to schematic plans provided (Supplemental Figures 1 to 5).

**Surgical planning**

All surgical plans were generated in Brainlab Elements software (Brainlab AG, Munich, Germany), using SpiraMed needle/catheter files in .stl format

**Catheter Deployment**

Catheter dimensions were determined using digital calipers to measure helix pitch and diameter, with corrections applied to each measured value to account for the catheter outside diameter (0.91mm).

**Agarose blocks and brain phantoms**

Agarose targets were prepared by dissolving the relevant weight of agarose powder (Sigma-Aldrich) in 200 mL of deionised water by heating to 100+ °C using a hotplate for 15 mins. Once dissolved, the solution was allowed to cool at room temperature for 15 mins before being poured into transparent acrylic sample containers or deformable molds, which were left at room temperature to solidify overnight.

**Therapy Delivery *in vitro* testing**

The SpiraMed device was first primed with a methylene blue solution at a concentration of 0.2 mg/mL in deionised water. Using a Lab Jack, the probe was slowly lowered into the agarose gel containers until a depth of 5.75 cm was reached. The SpiraMed coil was then deployed for 110 mm with a rotation rate of 5 deg/mm using the SpiraCell GUI. Once the deployment was complete a 25 µL bolus of the methylene blue solution was injected followed by a full retraction of the coil, during which a fluid dispense rate of 1.14, 2.05, or 2.95 µL/mm was utilised to deliver 150, 250, or 350 µL in total solution (including the bolus) to the agarose gel. After the probe was fully removed from the gel using the Lab Jack, the samples were imaged using a custom brain phantom monitoring and modelling platform previously published [1,2].

Straight-needle experiments were conducted using the SpiraMed coil tubing for comparison purposes. The tubing was primed with 0.2 mg/mL of methylene blue solution, before being inserted into agarose gel to a depth of 6.75 cm. Solution injection for single insertion was conducted by first injecting 50 µL at the maximum depth, followed by retracting the tubing 5 mm and injecting another 50 µL. The retraction and injection were repeated 3 more times for a final injection volume of 250 µL across 2 cm. Solution injection for 5 insertions was conducted by first injecting 10 µL at the maximum depth. A 5 mm retraction and 10 µL injection step was then repeated 4 times for a final injection volume of 50 µL across 2 cm per insertion. Insertion 2 and 3 were conducted by tilting the agarose gel container 4 degrees clockwise and counterclockwise and using the same entry point as the initial insertion. Insertion 4 and 5 followed the same process but with a tilt angle of 8 degrees. After the full delivery of the solution, and removal of the tubing, the samples were imaged using the custom brain phantom monitoring and modelling platform previously published ^1,2^.

**hESC Maintenance**

**Ventral midbrain dopaminergic progenitors and neurons**

Progenitors and neurons were differentiated from RC17 human embryonic stem cells using a hybrid of published ‘Lund’ and ‘Edinburgh’ protocols^3,4^.

**Human embryonic stem cells** from the RC17 cell line (Roslin Cells, hPSCreg number: RCe021-A) were maintained on Laminin-521 (1 μg/cm^2^, Biolamina) in StemMACS iPS-Brew XF media (Miltenyi Biotech) with media changes performed every day. Once cells reached a confluency between 70-90% they were washed with PBS (−Ca^2+^/−Mg^2+^, Thermo Fisher Scientific), passaged using EDTA (0.5mM, Thermo Fisher Scientific) and replated. For the first 24 hours following replating, StemMACS Y-27632 (10 μM, Miltenyi Biotec) was added to the medium; thereafter, it was omitted.

**Progenitor** differentiation batches were maintained at 37 centrigrade 5% CO₂, 21% O₂. On day 0 cells were dissociated with 0.5mM EDTA and seeded at 40,000 cells/cm2 on culture vessels coated with Laminin-111 (2 μg/cm2, Biolamina, LN-111). From day 0 to day 4, cells were differentiated in neural induction medium (NIM) which consisted of 50% DMEM/F12 (Thermo Scientific, 21331020), 50% Neurobasal Media (Thermo Scientific, A1371201), B27 supplement (without Vitamin A, 1:50, Thermo Scientific, 12587010), N2 supplement (1:100, Thermo Scientific, A1370701) and L-Glutamine (2 mM, Thermo Scientific, 25030081). Y-27632 (10 μM) was maintained in NIM for 48 hours after seeding. From day 0 to day 9, cultures received SB431542 (SB, 10 μM, Miltenyi, 130-106-543), LDN-193189 (LDN, 100ng/ml, Miltenyi Biotec, 130-106-540), Shh51 C24II (SHH, 600ng/ml, Miltenyi, 130-095-727) and CHIR99021 (CHIR, 1.3 μM, Miltenyi, 130-106- 539). From day 4 to day 11, medium was changed to neural patterning medium (NPM) comprising 50% DMEM/F12, 50% Neurobasal Media, B27 supplement (1:100), N2 supplement (1:200) and L-Glutamine (2 mM). From day 9 to day 16, FGF8b (100 ng/ml, Miltenyi, 130-095-740) and heparin (1 μg/ml, Sigma-Aldrich, H3149) were added. On day 11, cells were dissociated with Accutase (Thermo Scientific, A1110501) and replated to Laminin-111-coated (2 μg/cm2) plates at a density of 800,000 cells/cm2. From day 11, cells were fed with neural differentiation medium (NDM) which was made up of Neurobasal Media, B27 supplement (1:50), L-Glutamine (2 mM), and supplemented with L-Ascorbic acid (AA, 0.2 mM, Sigma-Aldrich, A4403-100MG), brain-derived neurotrophic factor (BDNF, 20 ng/ml, Miltenyi, 130- 096-286) and glial cell line-derived neurotrophic factor (GDNF, 10 ng/ml, R&D Systems, 212-GD-010), as well as FGF8b and heparin.

**Neurons** were produced by washing day 16 progenitors with PBS (−Ca^2+^/−Mg^2+^, Thermo Fisher Scientific), passaging using Accutase and replating on Laminin-111 (2 μg/cm^2^, Biolamina) at a density of 800,000 cells per cm^2^. These cells were maintained in NDM supplemented with StemMACS Y-27632 (from day 16-18 only) (10 μM, Miltenyi Biotec), BDNF (20 ng/ml, Miltenyi Biotec), GDNF (10ng/ml, Miltenyi Biotec), AA (0.2 mM, Sigma Aldrich), db-CAMP (0.5mM, Sigma Aldrich) and DAPT (1μM, Miltenyi Biotec) until day 45 with media changes carried out every 2-3 days. On day 45, the maturation of hESCs-derived mesDA progenitors into neurons is complete.

**Chimeric Antigen Receptor T-Cell generation**

Peripheral blood mononuclear cells (PBMCs) were isolated from healthy donor blood draws by density-gradient centrifugation using Lymphocyte Separation Medium (Corning, Cat. No. 25-072-CV) on day 0. T cells were subsequently isolated from PBMCs using the Pan T Cell Isolation Kit (Miltenyi Biotec, Cat. No. 130-096-535) according to the manufacturer’s instructions. Purified T cells were cultured in TexMACS Medium (Miltenyi Biotec, Cat. No. 130-097-196) supplemented with recombinant human IL-2 (100 ng/mL; Qkine, Cat. No. Qk089-0100) and IL-15 (5 ng/mL; Qkine, Cat. No. Qk097-0050), and activated with T Cell TransAct (Miltenyi Biotec, Cat. No. 130-111-160). Cells were maintained at 37°C in a humidified atmosphere containing 5% CO₂. On day 1, T Cell TransAct was removed, and the cells were maintained in TexMACS Medium supplemented with IL-2 (100 ng/mL) and IL-15 (5 ng/mL). On day 4, activated T cells were transduced with CAR-encoding lentiviral vectors ^5^ at a multiplicity of infection (MOI) of 5 using recombinant human fibronectin fragment (Takara Bio, Cat. No. T100A) according to the manufacturer’s instructions. Following transduction, CAR-T cells were cultured for an additional 5 days in TexMACS Medium supplemented with IL-2 (100 ng/mL) and IL-15 (5 ng/mL) before being used for subsequent experiments.

**SpiraMed biocompatibility testing**

The SpiraMed delivery catheter was preloaded with PBS by aspiration. **Dopaminergic progenitor** cells were harvested under sterile conditions and resuspended at 8 × 10⁶ cells/mL in 300ul flow media compositing of PBS (Sigma Aldrich,D8537-500ML) +10% KnockOutSerumReplacement (ThermoFisher 10828028 500ml). 330μl of the cell suspension was loaded into SpiraMed, by aspiration through the delivery needle tip as envisaged for intracerebral delivery. Following loading, cells were recovered from the device by flushing 330μl, and the recovered cell suspension was collected for viability analysis. An equivalent aliquot of the original cell suspension, prior to loading, was analysed in parallel to provide a baseline viability measurement. **CAR-T cells** were harvested under sterile conditions and resuspended in TexMACS Medium at a concentration of 1 × 10⁶ cells/mL, supplemented with recombinant human IL-2 and IL-15. For each sample, 300 µL of the cell suspension was loaded into the SpiraMed catheter, by aspiration through the delivery needle tip as envisaged for intracerebral delivery.  Cells collected following passage through the SpiraMed catheter were either analysed immediately or cultured in 48-well plates for analysis on the following day.

**Cell viability assessment**

Untreated control and SpiraMed-delivered samples were collected into microcentrifuge tubes. Cells were centrifuged at 300g for 5 mins, supernatant aspirated and resuspended in 500 uL flow media (BD Biosciences, catalogue no. 342003). Cells were then incubated with Hoechst 33342 (Invitrogen, catalogue no. H3570) and Propium Iodide (Invitrogen, catalogue no. P3566) or Incucyte® Cytotox Green Dye (Sartorius, 4633) for 10-30 minutes at manufacturer specified concentrations. Unstained, live-cell, and dead-cell controls were used to establish background fluorescence and define live/dead gating. The dead-cell control was generated by incubating with 0.1% triton X-100 (Fisher Scientific, cat. no. 10254640) for 10 minutes. Samples were analysed by flow cytometry using BD Fortessa X-20. A consistent acquisition threshold was applied across samples, with 10,000 cell events aquired per sample. BD FACSDivaTM was used to apply gating during acquisiton. Post sample-acquisition, further flow cytometry analysis was performed using the FlowJo11TM software. Cells were first identified using forward scatter and side scatter properties. Doublets and debris were excluded using singlet and scatter-based gates. Live and dead populations were then identified using the live/dead fluorescence channel, with gates defined using the unstained, live-control, and dead-control samples.

**Large animal testing**

Testing was performed at the South Australian Health and Medical Research Institute Large Animal Research and Imaging Facility. Male and female merino sheep aged ca 3 years were underwent surgery under the provisions of *SAM-25-018- Staged ovine model to determine the efficacy of a novel spiral brain cell delivery system*. Stage 1 comprised testing in 3 cadaveric heads. Stage 2 comprised live animal non-recovery testing under general anaesthesia.

**Anaesthesia -** 1000 mg of levetiracetam was administered intravenously pre-op. Each fasted sheep was anaesthetised with intravenous (jugular) ketamine (0.05 ml/kg) and diazepam (0.08 ml/kg) and a jugular catheter inserted. The animal was then intubated and mechanically ventilated using an oxygen (500 ml/min) and air (3.5 L/min) mix. General anaesthesia was maintained with a combination of inhalational isoflurane (1.5-2%) and intravenous ketamine (4 mg/kg/hr). An arterial line was placed in the artery of the ear to collect arterial blood gas samples and allow continuous measurement of blood pressure and calculation of cerebral perfusion pressure.

**Surgery -** MRI was obtained including T1/2 sequences. A Kopf Instruments 1630 Stereotactic Frame was attached. CT scan was obtained and fused with the T1/2 sequences of the MRI. Wool was removed from the scalp with clippers. Bilateral 3cm incisions were made over the frontal bone corresponding to pre-planned entry points. Bilateral frontal burr holes were placed. The dura was opened in cruciate fashion with a scalpel, the edges cauterised with bipolar forceps. A small corticectomy is made on the surface of the brain. On the right side SpiraMed was inserted into the Putamen with Stereotactic guidance. The SpiraMed catheter was then advanced through the striatum. Once the catheter was at maximum extension it was then retracted at a rate of 1mm per second whilst simultaneously injecting contrast. Following complete retraction the needle was withdrawn. On the Left side conventional stereotactic delivery was performed with division of the injection volume between three boluses for each trajectory, delivered interspersed with needle retraction steps. Finally the stereotactic frame was removed and Post-procedure MRI was performed.

**Sacrifice and processing-** Following completion of the MRI, all animals were humanely killed under general anaesthesia via carotid artery infusion with tris-saline Tris(hydroxymethyl) aminomethane and sodium chloride followed by jugular exsanguination. The neck was dissected to expose the carotid arteries, and infusion lines were inserted bilaterally into the lumen of the carotids and secured into place. Heparin (25000 IU in 5 mL 0.9% sodium chloride) followed by a 10 mL 0.9% sodium chloride flush was administered via jugular catheter to prevent coagulation. A perfusion pump wasattached, and at a pressure of 140 mmHg, sheep perfused with ~4 L of chilled Tris via the common carotid artery lines. Post-mortem the brain was removed and up to 10 coronal slices taken for fixation and histopathological analysis of tissue architecture.

**MRI volume segmentation-** T2-weighted MR images were imported into ITK-SNAP, an open-source software platform for the visualisation and segmentation of three-dimensional medical imaging data (Yushkevich et al., 2006). ITK-SNAP enables manual voxel-based labelling of structures on two-dimensional image slices and subsequent three-dimensional visualisation and volumetric analysis of the resulting segmentation.

Regions of corresponding to the intracerebral distribution of injected dye were manually delineated on consecutive image slices using the manual segmentation tools in ITK-SNAP. Voxels within the boundaries of the observed dye-associated signal change were assigned to the region of interest (ROI), producing a three-dimensional segmentation mask representing the spatial distribution of the injectate. Segmentations were reviewed in the orthogonal imaging planes to ensure spatial consistency and appropriate delineation of the signal abnormality. The resulting three-dimensional ROIs were used to calculate the volume of dye distribution.

**Cadaveric human testing**

Cadaveric human heads were prepared in the Cambridge University Hospitals Microsurgical Skills, and the Leksell G stereotactic frame was applied. A CT volume scan was performed to enable Brainlab software planning. Burrholes were drilled, and the Generation 1 SpiraMed prototype inserted and deployed. 1000ul of Omnipaque-240 CT contrast diluted 1:20 was administered to each hemisphere synchronized with catheter retraction. A further CT scan was performed to visualise contrast delivery.

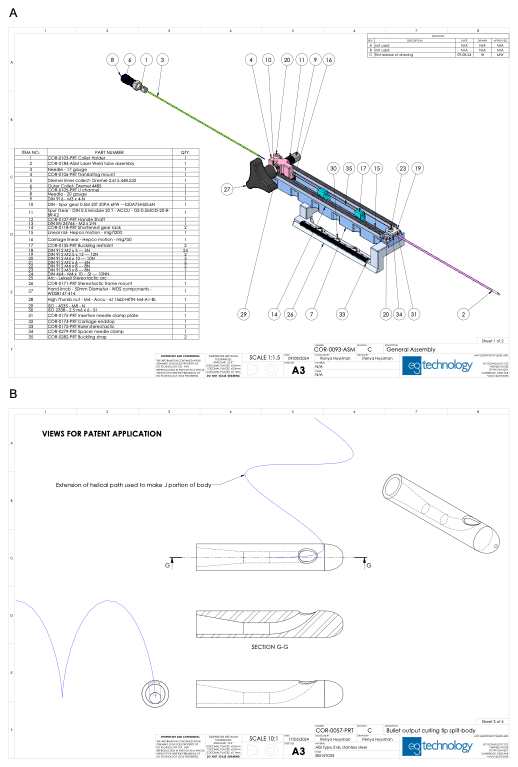

**Supplemental Figure 1 A**: SpiraMed Generation 1 mechanically-actuated prototype schematic **B** ‘Twisted-J’ needle tip for helix deployment.

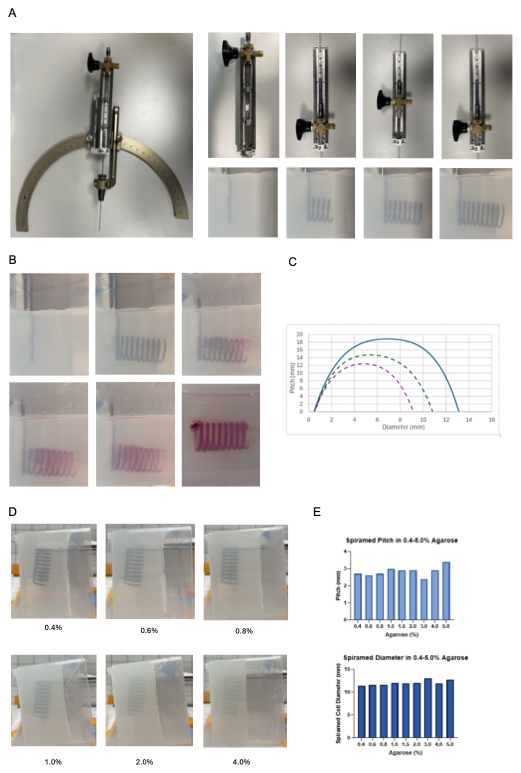

**Supplemental Figure 2 A**: SpiraMed Generation 1 mechanically-actuated prototype - mounted on Leksell G stereotactic arc (left panel) and demonstrating helix deployment into agarose (right panels) **B** Helix deployment and delivery of red dye payload **C** Helix dimensions for tip radius 6mm (blue), 4.5mm (green), 3mm (purple) **D,E** Example deployments and measured helix pitch and diameter, in agarose blocks composition as indicated.

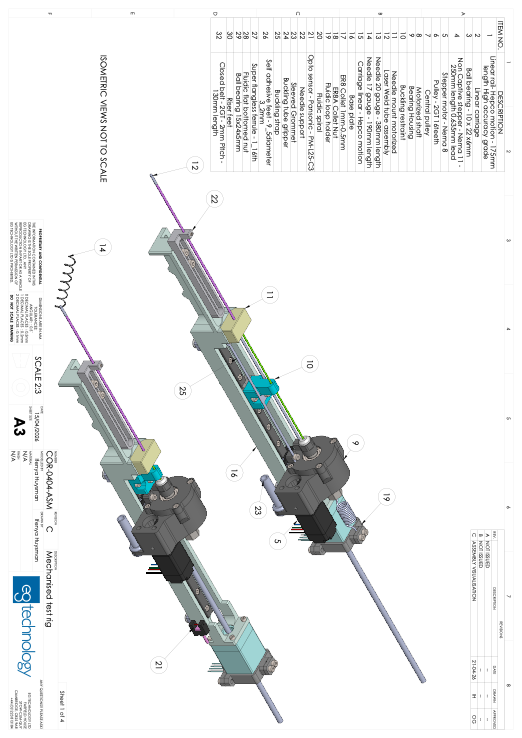

**Supplemental Figure 3:** SpiraMed Generation 2 electronically actuated prototype schematic

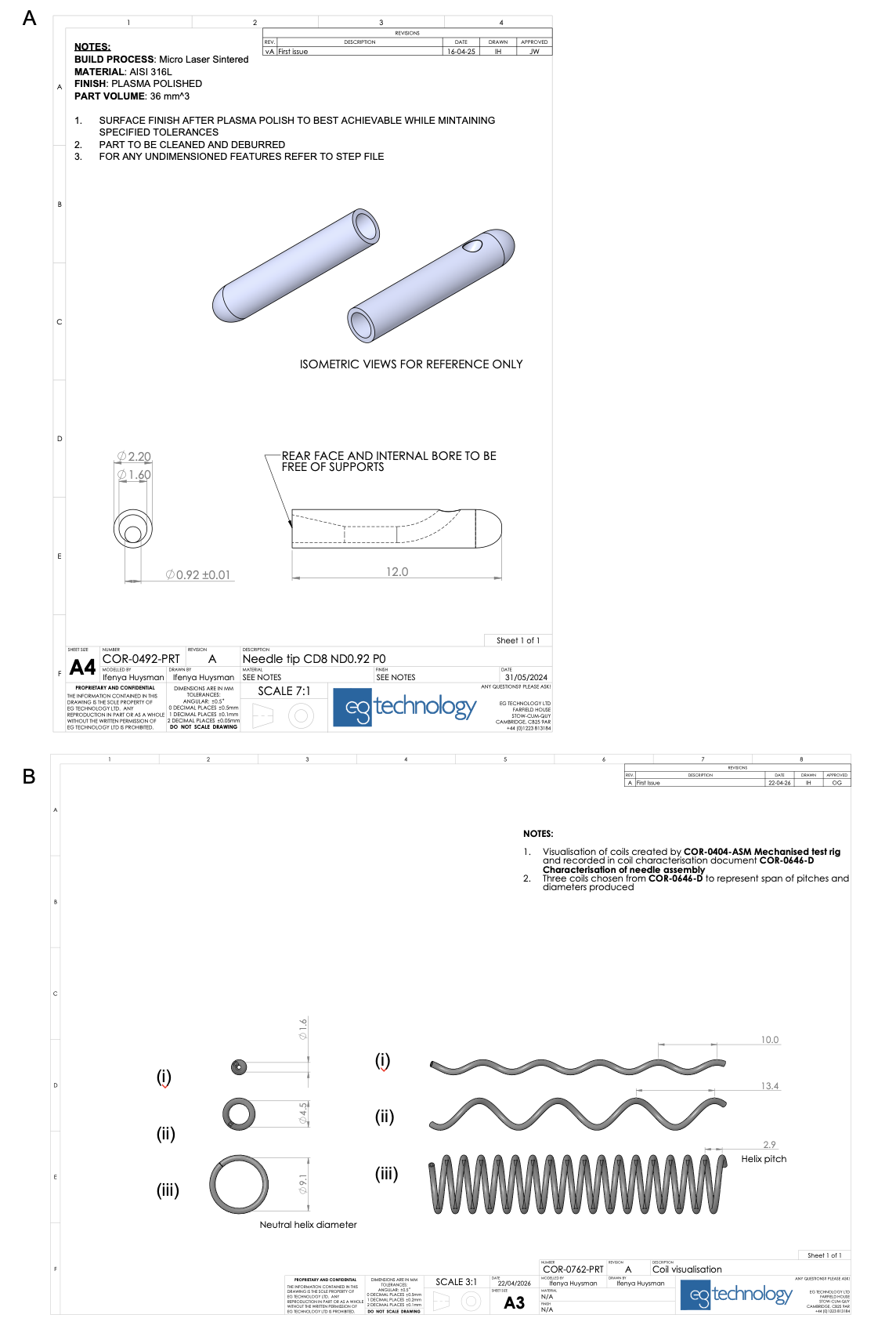

**Supplemental Figure 4:**

**A** SpiraMed Generation 2 prototype ‘unmodified’ J-shape needle tip **B** Example helices generated using tip radius **r** _tip_=4mm and catheter rotation rate (i) 35°/mm (d=1.6mm; p = 10.0mm) (ii) 15°/mm (d=4.5mm; p = 13.4mm) (iii) 1°/mm (d=9.1mm; p = 2.9mm).

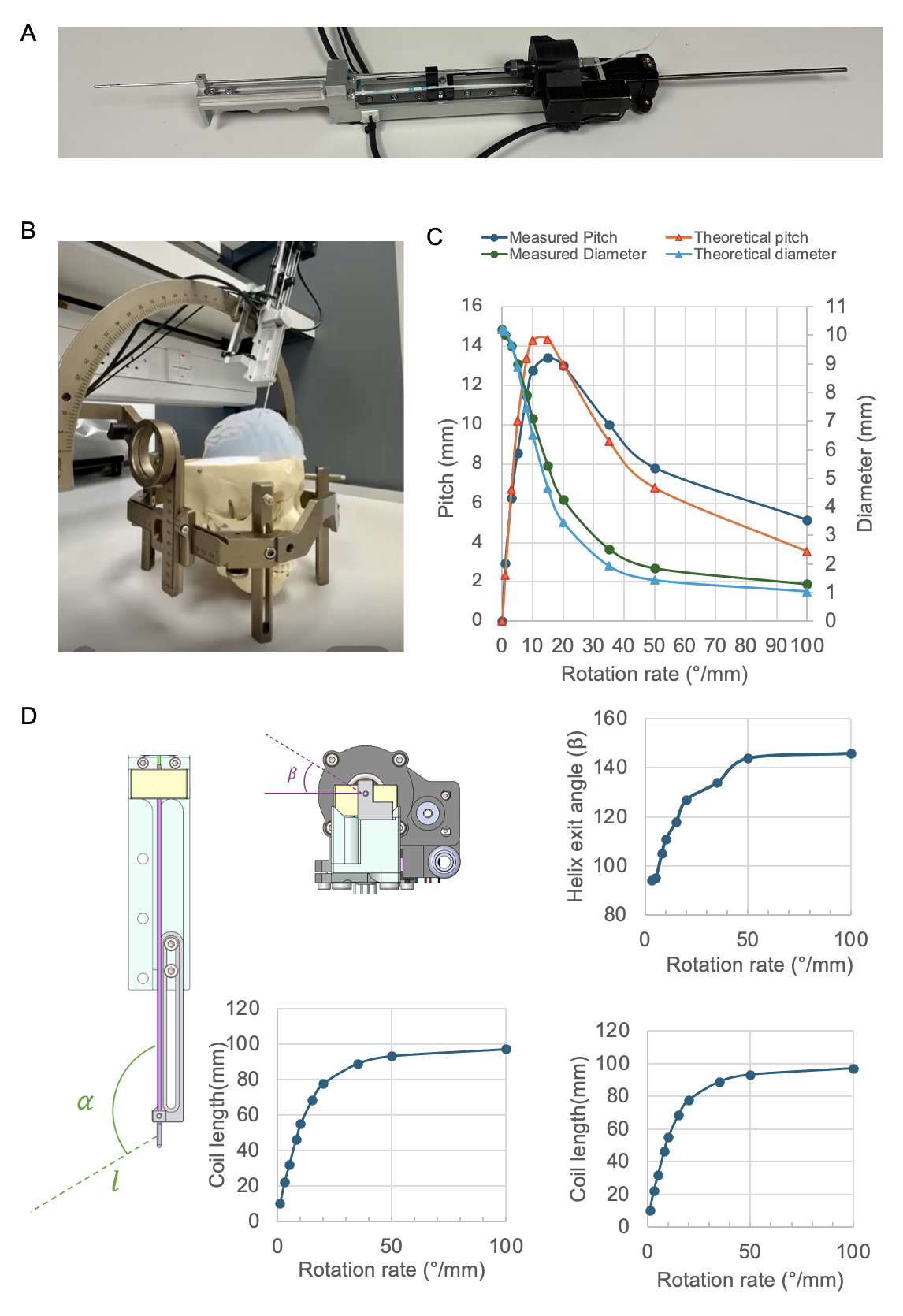

**Supplemental Figure 5: A,B** SpiraMed Generation 2 electronically actuated prototype benchtop and example Leksell G frame/ agarose brain phantom deployments **C** Measured helix pitch and diameter compared to Frennet-Serret framework predictions (rtip=4mm). **D** Additional predicted helix deployment parameters (rtip=4mm) for incorporation in future surgical planning software implementations.

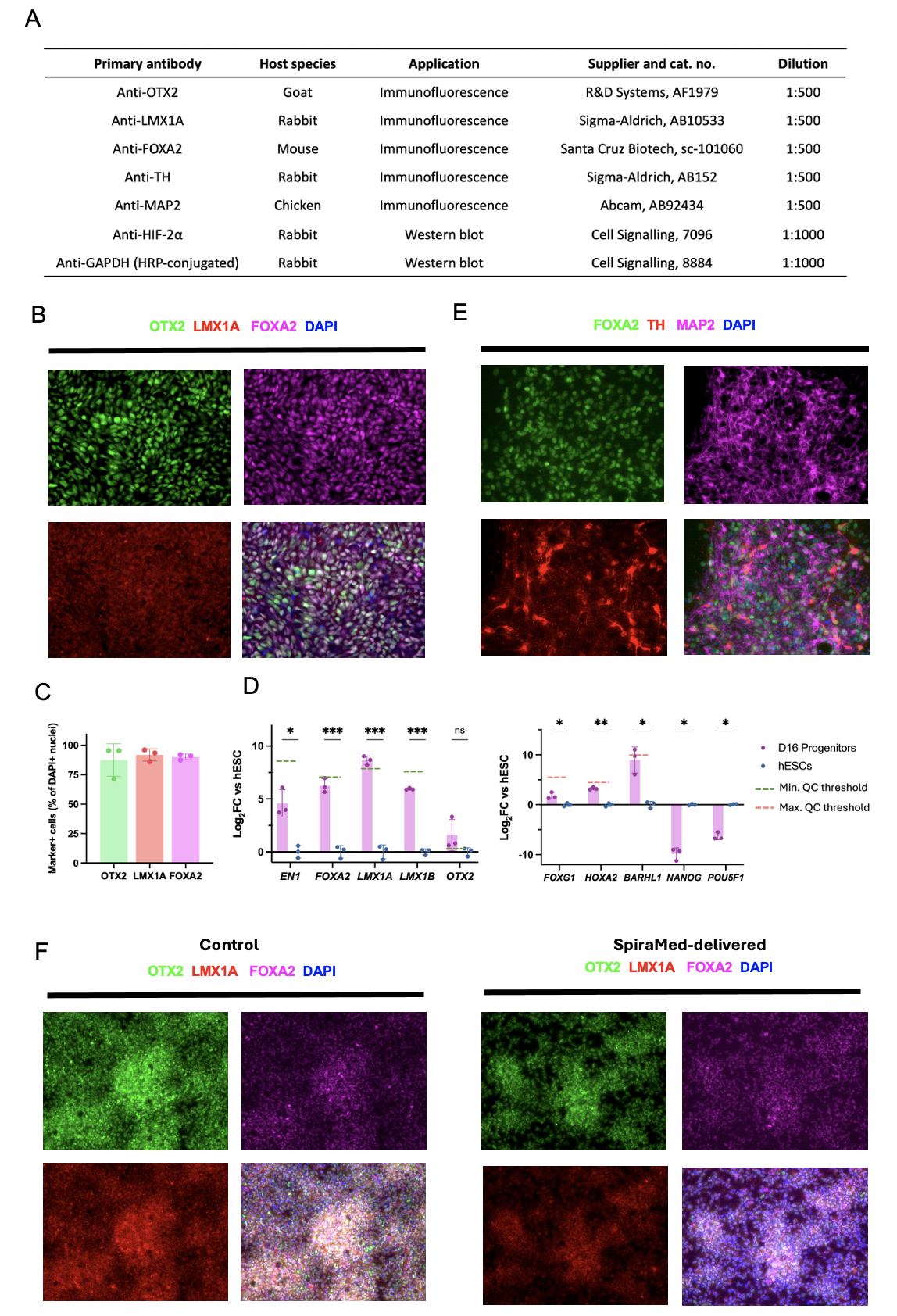

**Supplemental Figure 6: A** Primary antibodies used for vmDA progenitor cell characterisation. **B, C** Immunofluorescence and **D** qPCR quantifying Day 16 vmDA progenitor marker expression **E** Day 60 vmDA neuron marker expression **F** Immunofluorescence of vmDA progenitors - control (left panels) and SpiraMed-delivered (right panels).

### Supplemental References

1. Rognin, E., Willis-Fox, N. & Daly, R. Quantitative monitoring and modelling of retrodialysis drug delivery in a brain phantom. *Sci. Rep.* **13**, 1900 (2023).

2. Naegele, T. E. *et al.* Redox Flow Iontophoresis for Continuous Drug Delivery. *Adv. Mater. Technol.* **9**, (2024).

3. Nolbrant, S., Heuer, A., Parmar, M. & Kirkeby, A. Generation of high-purity human ventral midbrain dopaminergic progenitors for in vitro maturation and intracerebral transplantation. *Nat Protoc* **12**, 1962–1979 (2017).

4. Kunath, T. *et al.* Midbrain dopaminergic differentiation of human pluripotent stem cells v1. <https://doi.org/10.17504/protocols.io.bddpi25n> (2020) doi:10.17504/protocols.io.bddpi25n.

5. Tang, X. *et al.* B7-H3 as a Novel CAR-T Therapeutic Target for Glioblastoma. *Mol. Ther. - Oncolytics* **14**, 279–287 (2019).
