## Appendix 1 - Frennet Serret Framework for "SpiraMed – A Stereotactic Helix-based Therapy Delivery system for the Human Brain"

### Introduction

The coil-forming process is interpreted through the Frenet–Serret framework, which provides a geometric description of how curvature and torsion shape 3D curves. This framework is applied specifically to the operation of the device described in this work.

A set of simplified equations derived from the Frenet–Serret formalism are introduced to model the coil formation process. These equations are validated against experimental data, demonstrating their accuracy in predicting the resulting coil geometry.

The machine layout used in this process is believed to be novel, offering a unique approach to controlled 3D curve formation. Additionally, this work references existing techniques that also utilize the Frenet–Serret framework as a foundational tool for generating 3D curves, situating the current method within the broader context of established literature.

### Generation 1 Prototype Implementation

Generation 1 prototypes create helical coils by advancing a straight delivery needle through a helically curved inner path at the distal tip of an outer guide needle, plastically deforming it into a helical set. The helical guide path in the outer guide needle has closely matched inner diameter to that of the delivery needle. The guide needle at its proximal end is coupled to a carriage that is hand driven by user, providing manual advancing and retraction. The delivery needle terminated with a luer connection, onto which a manual syringe was attached. Limitations of Gen 1 devices came from inability to form coils with large pitch and small diameters. A matrix of guide needles were produced with varying internal pitches and diameter and the resulting delivery coils measured. For a given internal curve diameter increasing internal curve pitch resulted in minor change to output coil pitch. Thus operation flexibility of devices was limited to producing a narrow range of output coil pitches.

### Generation 2 Prototype Implementation

The coil formation technique used by the Gen 2 device is based on the combined linear advancement and axial rotation of an initially straight needle through a curved path. This curved path is planar and has a fixed radius, imposing a repeatable bending profile on the needle as it passes through the curve. On exiting the tip, the needle retains a deformed shape resulting from both plastic deformation and elastic recovery. During coil generation, the combination of needle axial and needle linear advancement produces a three-dimensional helical form. Experimental observations showed that, for a given tip curve radius, all resulting coils share an approximately constant curvature, while torsion is governed by the applied rotation rate. Rotation rate can be treated as approximately equal to torsion, enabling prediction of coil pitch and diameter from the imposed rotation and the effective curvature associated with the tip.

### Frenet-Serret frame and equations background

The Frenet–Serret formalism describes the geometry of smooth curves in three-dimensional space by constructing a moving local coordinate system along the curve. This system consists of three mutually orthogonal unit vectors that evolve as the curve progresses.

1. **Tangent Vector (T)**

The first vector is the tangent vector, which points in the direction of motion along the curve. For a curve parameterized by arc length the tangent vector is defined as the normalized derivative of the position vector r(*t*):

## 𝑟^′^(𝑡) 𝑇(𝑡) = ‖𝑟_′_(𝑡)‖

*Equation 1*

**2. Normal Vector (N)**

The second vector is the principal normal vector, which indicates the direction in which the curve is turning. It points to the instantaneous centre of the curve. It is defined as the normalized derivative of the tangent vector

## 𝑇^′^(𝑡) 𝑁(𝑡) = ‖𝑇_′_(𝑡)‖

*Equation 2*

3. **Binormal Vector (B)**

The third vector is the binormal vector, defined as the cross product of the tangent and normal vectors. It is orthogonal to both and completes the right-handed orthonormal basis:

𝐵(𝑡) = 𝑇(𝑡) × 𝑁(𝑡)

*Equation 3*

Together, these vectors form the Frenet–Serret frame (T,N,B), which moves smoothly along the curve. In the case of a helical curve, for example, the tangent vector is shown in purple, the normal vector in green, and the binormal vector in black.

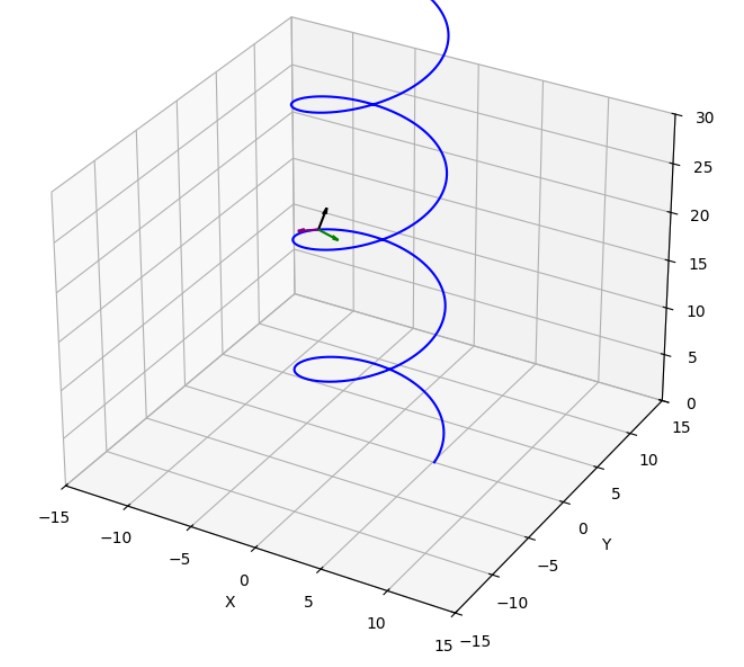

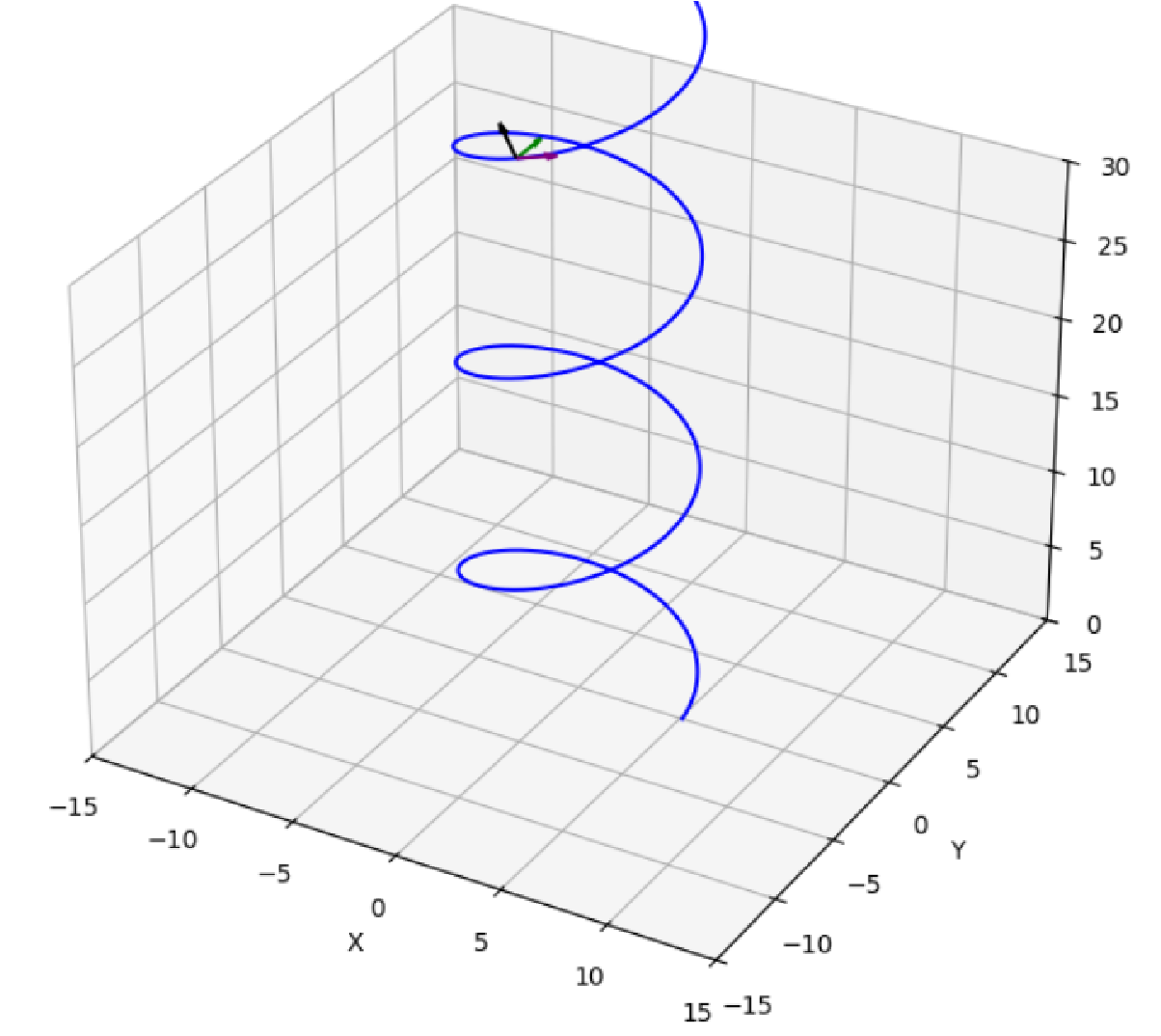

The Frenet–Serret formula describes curvature as the rate at which the tangent vector changes direction along a smooth curve. Curvature is the magnitude of this change, denoted by 𝜅.

The relationship is expressed as:

## 𝑑𝑇 𝑑𝑠

= 𝜅𝑁

*Equation 4*

A visualisation of curvature is the osculating circle, this is a circle that contacts point r(t) with equal tangent and curvature to the curve. It has radius, 𝑟_𝑜𝑠𝑐_

1

𝑟_𝑜𝑠𝑐_ = 𝜅(𝑡) (1)

*Equation 5*

The osculating circle of a helix is shown below as it progresses along the curve.

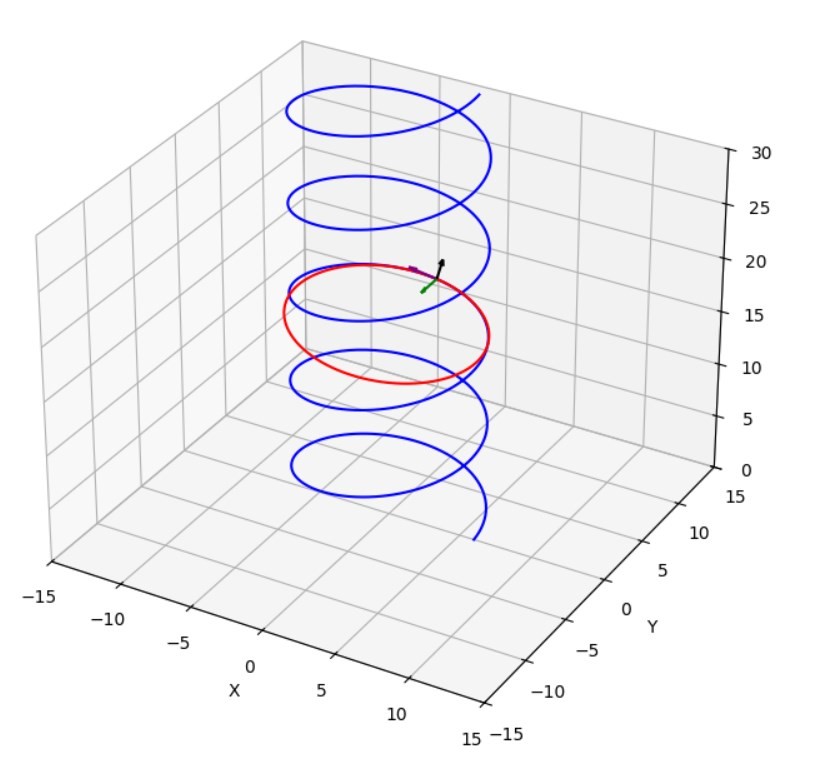

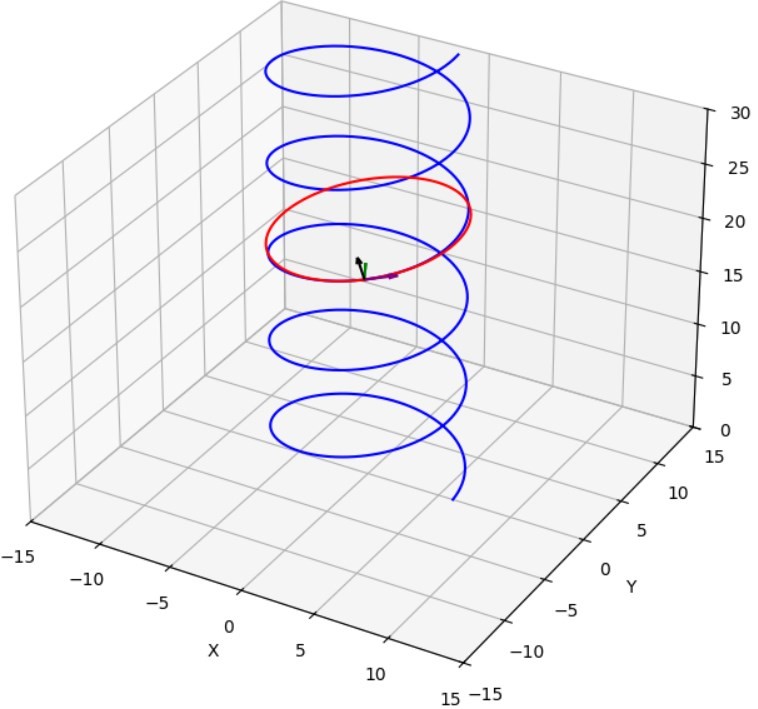

At any instant the plane this circle sits on is the osculating plane, defined by the vectors T and N.

Torsion *τ* describes how this osculating plane rotates as one moves along the curve. Specifically, it measures the rate of change of the binormal vector 𝐵, which is perpendicular to the osculating plane. This is described by the Frenet-Serret formula:

𝑑𝐵

= −𝜏𝑁

## 𝑑𝑠

*Equation 6*

This equation shows that the binormal vector changes in the direction opposite to the normal vector, and the magnitude of this change is the torsion. A nonzero torsion indicates that the curve is twisting out of the osculating plane, giving it a three-dimensional character.

The final equation in the Frenet–Serret formulas describes how the normal vector 𝑁 changes along the curve. This change is influenced by both the curvature and the torsion of the curve.

## 𝑑𝑁 𝑑𝑠

= −𝜅𝑇 + 𝜏𝐵

*Equation 7*

### Helix curvature and torsion

For a constant helix, the curvature 𝜅 depends on the radius of the helix and the spacing between its coils. Let:

- *r* be the radius of the helix,
- 𝑝 be the vertical distance between successive turns; pitch
- 𝑐 be the vertical rise per radian

𝑝 𝑐 =

2𝜋

*Equation 8*

## 𝑟

𝜅 = 𝑟_2_ + 𝑐2 (1) (2)

*Equation 9*

While torsion of a helix is described as

## 𝑐

𝜏 = 𝑟_2_ + 𝑐2 (1) (2)

*Equation 10*

This leads to the following being true:

## 𝜅 𝑟 𝜏 𝑐

=

*Equation 11*

Equations for torsion and curvature can be re-arranged as below:

## 𝜏

𝑐 = 𝜏2 + 𝜅2

*Equation 12*

## 𝜅

𝑟 = 𝜏2 + 𝜅2

*Equation 13*

### Coil formation

The 20Gauge needle is advanced through a planar curve in the needle tip, curve is planar as it is 2D in nature. It has constant radius 𝑟_𝑡𝑖𝑝_.

The curvature of the needle tip internal curve is then described as 𝜅_𝑡𝑖𝑝_

1

𝜅_𝑡𝑖𝑝_ =

𝑟𝑡𝑖𝑝

*Equation 14*

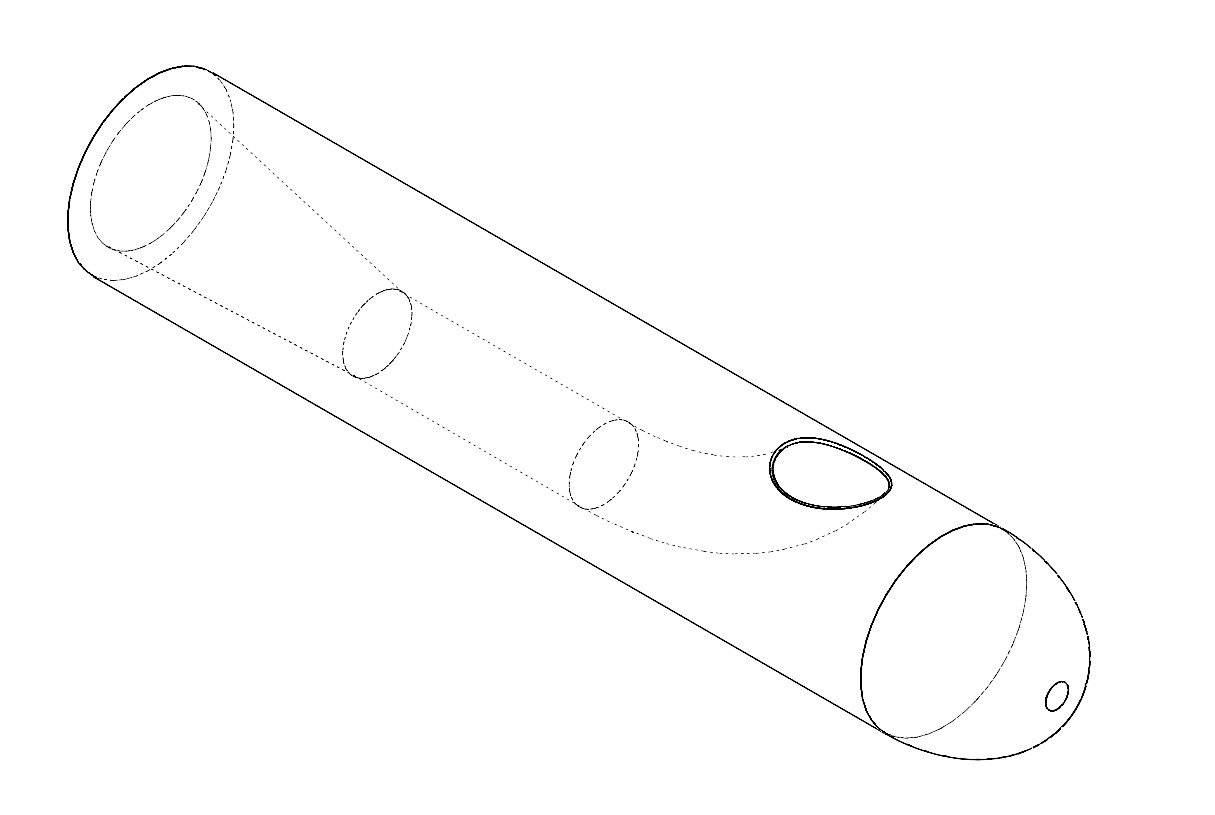

A needle advanced through the curve without axial rotation will adopt radius 𝑟 upon exit, with

𝑟 ≥ 𝑟_𝑡𝑖𝑝_

The needle undergoes both plastic and elastic deformation as it goes through the curve, the elastic component of this deformation is recovered upon exit of the curve; springback.

Springback prediction remains a topic of active academic interest, the following papers make strides in its prediction. (3) (4) (5) (6) (7)

The described technique did not attempt to predict springback, nor were advancements in springback understanding made over existing cited research.

The coil forming technique involves advancing needle through this curve with an added axial rotation. The needle thus experiences both linear and rotational motion as it transits into the curve. The rotation rate, 𝛼 is given in units (rad/mm), it is imparted to the needle by the rotational axis of the device.

For a given needle tip it was experimentally observed that all coils have equal curvature. A graph of measured curvature against rotation rate demonstrates this, noise in data primarily from challenge of physically measuring the coil dimension of pitch. Equation 12 utilised to determine curvature of the coils from the obtainable measurements of diameter and pitch.

0

0.02

0.04

0.06

0.08

0.1

0.12

0.14

0.16

0.18

0.2

0

0.05

0.1

0.15

0.2

0.25

0.3

Measured curvature (

𝜅

)

Rotation rate,

𝛼

(

rad/mm

)

All coils formed from a single tip share the same curvature, therefore for a given tip a single compensation factor 𝑓 relates coil curvature to the tip curvature.

𝜅 = 𝜅_𝑡𝑖𝑝_ 𝑓

*Equation 15*

Empirical measurement of coil curvature most accurate when 𝜏 = 0, as this produces a 2D curve, therefore Equation 9 reduces to being only in terms of *r.*

1

𝜅 =

𝑟

*Equation 16*

This can then be used to calculate compensation factor 𝑓

𝑟𝑡𝑖𝑝

𝑓 =

𝑟

*Equation 17*

In an entirely plastic situation 𝑓 would equal 1, as 𝑟_𝑡𝑖𝑝_ = 𝑟 . While in an entirely elastic situation 𝑓 would equal 0, as the needle would emerge from the coil undeformed 𝑟 = ∞.

Springback is encapsulated within the term 𝑓, while in this technique 𝑓 is experimentally determined, from the cited research it is feasible that a model accounting for material and geometric parameters could result in predictive form of it. This could capture the relationship between 𝑟_𝑡𝑖𝑝_ and 𝑓 allowing for predictions with alternative tip radiuses.

Rotation rate is observed as being approximately as equal to the torsion, a graph of rotation rate against measured torsion from coils produced shows this. Equation 10 used to derive torsion from physical measurements of coil pitch and diameter. Therefore rotation rate 𝛼 (rad/mm) can be equated to 𝜏

𝜏 = 𝛼

*Equation*

*18*

0

0.05

0.1

0.15

0.2

0.25

0.3

0

0.05

0.1

0.15

0.2

0.25

0.3

Measured torsion (

𝜏

)

Rotation rate,

𝛼

(

rad/mm

)

Equation 19 and Equation 20 are the critical equations that describe the relation that pitch and diameter (𝑑) have to rotation rate and the curvature of the needle tip.

2𝜅	2𝜅_𝑡𝑖𝑝_ 𝑓 𝑑 = 𝜅2 + 𝜏2 = (𝜅𝑡𝑖𝑝 𝑓)2 + 𝛼2

*Equation 19*

#### 2𝜋𝜏 2𝜋𝛼 𝑝 = 𝜅2 + 𝜏2 = (𝜅𝑡𝑖𝑝 𝑓)2 + 𝛼2

*Equation 20*

Values of pitch and diameter can be plotted against each other to yield the pitch/diameter curves associated with that tip. Pitch and diameter relationship shown for tip radius that results in curvature of coil 𝜅 of 0.082 and 0.057

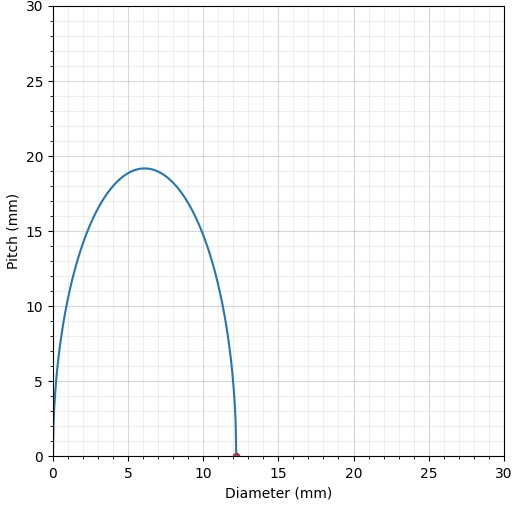

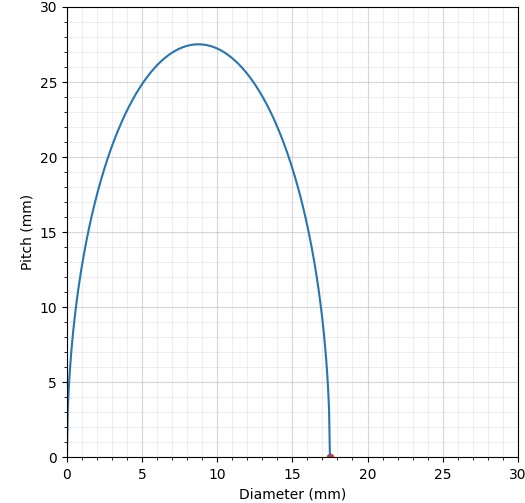

A comparison between the theoretical framework established above and measured values shows the high alignment of the two.

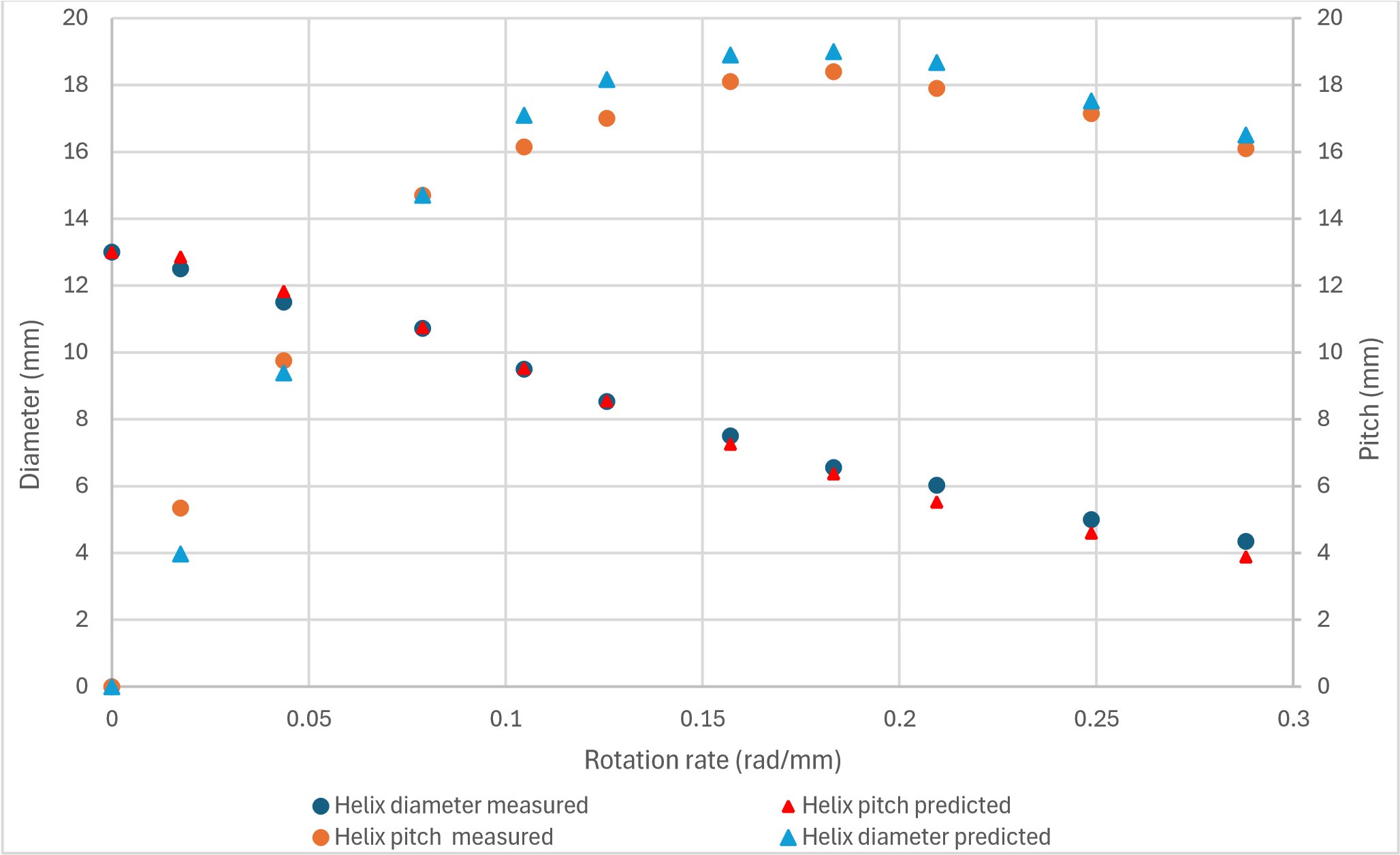

A simplification at the basis of this derivation is that the diameter of the helix is larger than needle diameter, in the framework above the needle is modelled as just a line; zero thickness. Closer alignment between theory and practical could be expected from a derivation that accounts for true wire geometry.

### Summary

The coil-forming technique can be effectively described using the Frenet–Serret framework, which characterizes the geometry of space curves through curvature and torsion.

In this process, a straight needle is transformed into a 3D curve by imparting:

- Curvature: Introduced by guiding the needle through the needle tip, which imposes a consistent bending profile. This sets the curvature of the resulting coil.
- Torsion: Applied by rotating the needle about its longitudinal axis. This rotation causes the curve to twist out of the osculating plane, introducing torsion into the coil.

The result is a helical coil with:

- Constant curvature, determined by the fixed curve of the needle tip
- Modifiable torsion, controlled by selecting the rotation rate of the rotary axis

This technique allows for precise and predictable control of coil geometry

### References

1. **Helix. *Wolfram.* [Online] https://mathworld.wolfram.com/Helix.html.**
2. **Wikipedia. *Frenet Serret Formula.* [Online] https://en.wikipedia.org/wiki/Frenet%E2%80%93Serret_formulas.**
3. ***Precise Pipe-Bending by 3-RPSR Parallel Mechanism Considering the Effect of Springback and Dies Clearances.* Kawasumi, Shohei and Takeda, Yukio and Matsuura, Daisuke. s.l. : Transactions of the JSME (in Japanese), 2014. 10.1299/transjsme.2014trans0343.**
4. ***A review on flexibility of free bending forming technology for manufacturing thin-walled complex-shaped metallic tubes.* Ali Abd El-Aty, Xunzhong Guo, Myoung-Gyu Lee, Jie Tao, Yong Hou, Shenghan Hu, Tao Li, Cong Wu, Qiucheng Yang,. 2, s.l. : International Journal of Lightweight Materials and Manufacture, Vol. 6. ISSN 2588-8404.**
5. ***Springback prediction model and its compensation method for the variable curvature metal tube***

***bending forming.* Zhang, S., Fu, M., Wang, Z. s.l. : The International Journal of Advanced**

**Manufacturing Technology , 2021, Vol. 112. https://doi.org/10.1007/s00170-020-06506-0.**

6. ***Spatial variable curvature metallic tube bending springback numerical approximation prediction and compensation method considering cross-section distortion defect.* Wang, Zili & Lin, Yaochen & Qiu, Lemiao & Zhang, Shuyou & Fang, Dingyu & He, Ci & Wang, Le. s.l. : The International Journal of Advanced Manufacturing Technology, 2022. 10.1007/s00170-021-08051-w. .**
7. ***Numerical simulation and experimental study on mechanism and characteristics of tube freebending forming process.* Guo, Xunzhong and Xiong, Hao. s.l. : Procedia Manufacturing, Vol. 15.**

**10.1016/j.promfg.2018.07.179.**

8. **Osculating circle. [Online] https://mathworld.wolfram.com/OsculatingCircle.html.**
